## Supplementary Information for "Functional Protein Dynamics in a Crystal"

### 1 Prior MD simulation studies of protein crystals

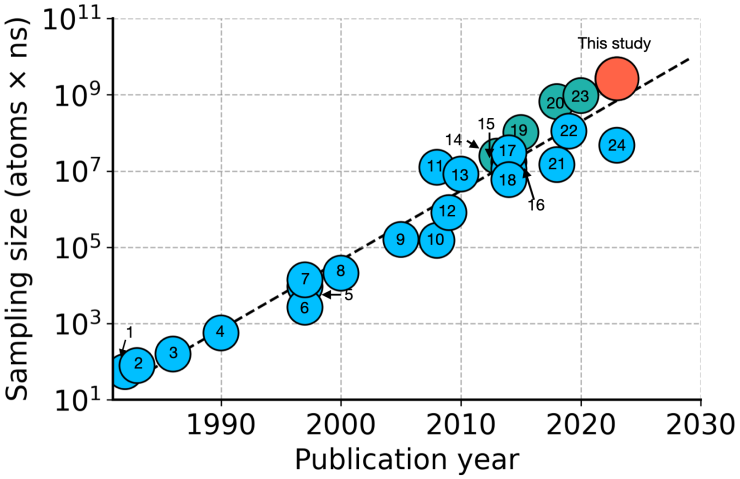

**Supplementary Figure** 1: **"Moore's Law" for MD simulations of protein crystals.** The total sampling (the total number of atoms in a simulation system × the simulation time in $ns$) as a function of the publication year is shown in a semi-log plot. Data points are enumerated according to the reference number (Supplementary References 1 to 24), with details for each study provided in Supplementary Table 1. Green and orange points represent the studies where the crystal systems were simulated for more than 1 $\mu$s. The dashed line represents a linear fit with a slope of $\approx$ 1/5 years$^{-1}$, which implies that the accessible sampling increases tenfold every five years due to advances in computational performance.

**Supplementary Table** 1: **Prior MD simulation studies of protein crystals.** For studies in which multiple systems were simulated, the system with the largest total sampling (simulation time × system size) is provided. When the total number of atoms was not provided, the system size was inferred from the information provided in the study to reconstruct the crystal cell. These inferred system sizes are shown in parentheses. The data in this table was used to make Supplementary Fig. 1. Note that this list of crystal MD studies is not exhaustive and is only meant to demonstrate the general trend in accessible sampling over time.

| Year | System | Simulation time (ns) | System size in atoms | Force field | Supercell / Unit cell | Ref. |
|---|---|---|---|---|---|---|
| 1982 | Bovine Pancreatic Trypsin Inhibitor | 0.025 | 2190 | Described in ref. | Unit cell | [1] |
| 1983 | Bovine Pancreatic Trypsin Inhibitor | 0.020 | 3948 | Described in ref. | Unit cell | [2] |
| 1986 | Bovine Pancreatic Trypsin Inhibitor | 0.040 | 3952 | Described in ref. | Unit cell | [3] |
| 1990 | Protease A (Streptomyces griseus) | 0.060 | 9427 | Full Valence Force Field (Hagler 1985) | Asymmetric Unit | [4] |
| 1997 | Crambin | 5.1 | 1830 | Amber ff94 | Unit cell | [5] |
| 1997 | Bovine Pancreatic Trypsin Inhibitor Type I | 0.25 | 10670 | Amber ff94, CHARMM ver. 23 | Supercell (1x2x1) | [6] |
| 1997 | Hen Egg-White Lysozyme | 1 | (13980) | CHARMM22 | Unit cell | [7] |
| 2000 | Hen Egg-White Lysozyme (1AKI) | 2 | 10469 | GROMOS96 ver.43A1 | Unit cell | [8] |
| 2005 | Staphylococcal nuclease (2SNS) | 10 | 15993 | CHARMM22 | Unit cell | [9] |
| 2008 | Staphylococcal nuclease (1STN) | 10 | 15108 | Amber ff99 | Unit cell | [10] |
| 2008 | Biotin-Liganded Streptavidin (1MK5) | 250 | (50000) | Amber ff99SB | Unit cell | [11] |
| 2009 | Hen Egg-White Lysozyme (1HEL) | 10 | (81000) | OPLS-AA, Amber03, GROMOS96 | Supercell (1x1x2) | [12] |

| Year | System | Simulation time (ns) | System size in atoms | Force Field | Supercell / Unit cell | Cit. |
|---|---|---|---|---|---|---|
| 2010 | Toxin protein II (Androctonus australis, 1AHO) | 100 | 83000 | Amber (ff99SB, ff03), CHARMM22, and OPLS | Supercell (2x2x3) | [13] |
| 2013 | Synthetic fav8 decapeptide | 2400 | 10224 | Amber ff99SB | Supercell (4x3x3) | [14] |
| 2014 | Staphylococcal nuclease (1STN) | 1100 | 15421 | OPLS-AA | Unit cell | [15] |
| 2014 | Ubiquitin (3ONS) | 200 | 56240 | Amber ff99SB*-ILDN | Supercell (2x2x1) | [16] |
| 2014 | Toxin protein II (Androctonus australis, 1AHO) | 250 | (120000) | Polarized Protein-Specific Charge, Amber ff99SB | Supercell (3x3x3) | [17] |
| 2014 | Villin headpiece domain (2F4K) | 50 | 118752 | GROMOS 45A3 | Supercell (3x3x3) | [18] |
| 2015 | Hen Egg-White Lysozyme (4LZT) | 3000 | (34000) | Amber (ff99SB, ff14ipq, ff14SB) and CHARMM36 | Supercell (2x2x3) | [19] |
| 2018 | Ternary Complex of Staphylococcal Nuclease (1SNC) | 5100 | 129462 | CHARMM27 | Supercell (2x2x2) | [20] |
| 2018 | Hen Egg-White Lysozyme (2VB1) | 250 | (58900) | Polarized Protein-Specific Charge, Amber ff99SB | Supercell (3x3x3) | [21] |
| 2019 | Staphylococcal Nuclease (4WOR) | 600 | (185000) | CHARMM27, Amber ff14SB | Supercell (2x2x2) | [22] |
| 2020 | Hen Egg-White Lysozyme (4LZT) | 1000 | (950000) | Amber ff14SB | Supercell (7x7x7) | [23] |
| 2023 | Catalytic subunit of mouse protein kinase A (7UJX) | 100 | (473000) | Amber ff14SB | Supercell (2x2x2) | [24] |
| 2024 | The second PDZ domain of LNX2 protein (5E11) | 10000 | 261852 | CHARMM36m, Amber ff14SB, Amber ff94 | Supercell (3x3x3) | This study |

### 2 Supplementary Results

#### 2.1 Fraction of native contacts, Q

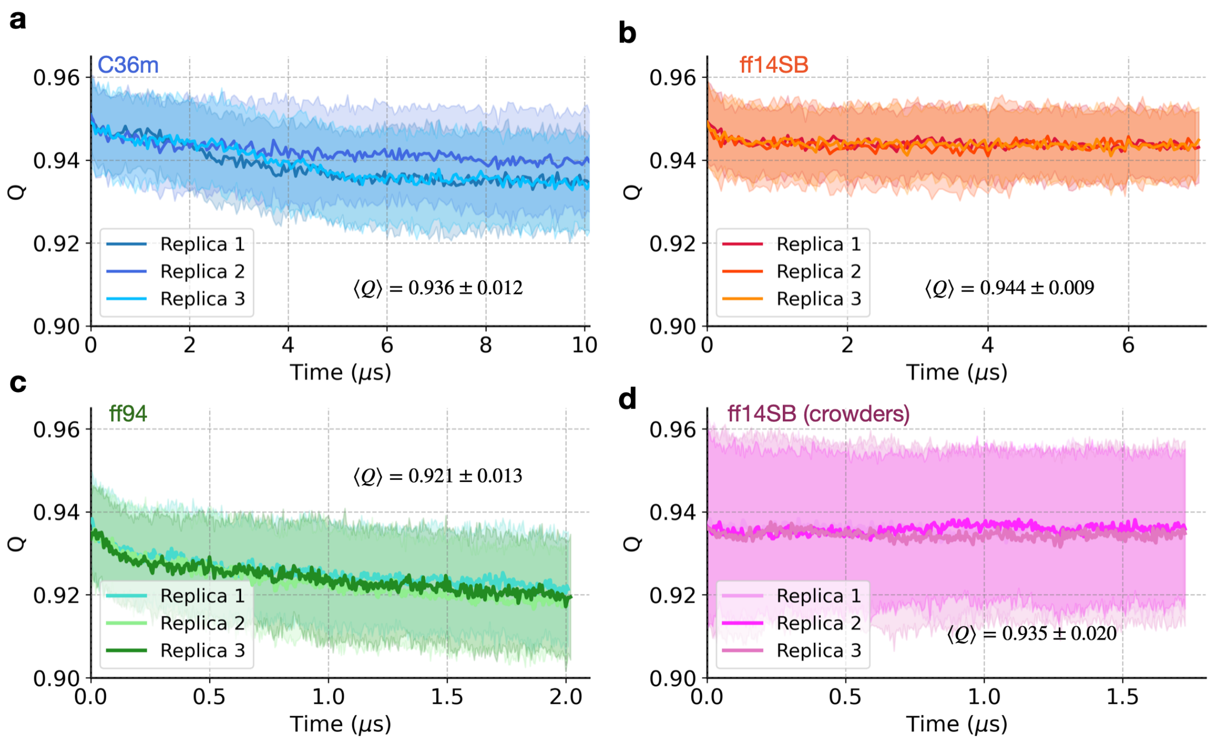

**Supplementary Figure** 2: **Fraction of native contacts, Q.** The average fraction of native (intra-protein) contacts, Q, in the supercell simulations is shown for the simplified crystal environment using force fields (**a**) C36m, (**b**) ff14SB, and (**c**) ff94, as well as (**d**) ff14SB with crowders. The standard deviation ($n = 108$) for each replica is represented by a shaded envelope. Each plot shows $\langle Q \rangle$ ($\pm$ standard error for $n = 3$ replicas) computed for the last 1 $\mu$s of simulation. Note that the simulations with (**c**) ff94, and (**d**) ff14SB with crowders have not reached equilibrium. For further details regarding the method of counting native contacts, refer to SI section 3.4.

#### 2.2 Convergence of the atomic covariance matrix

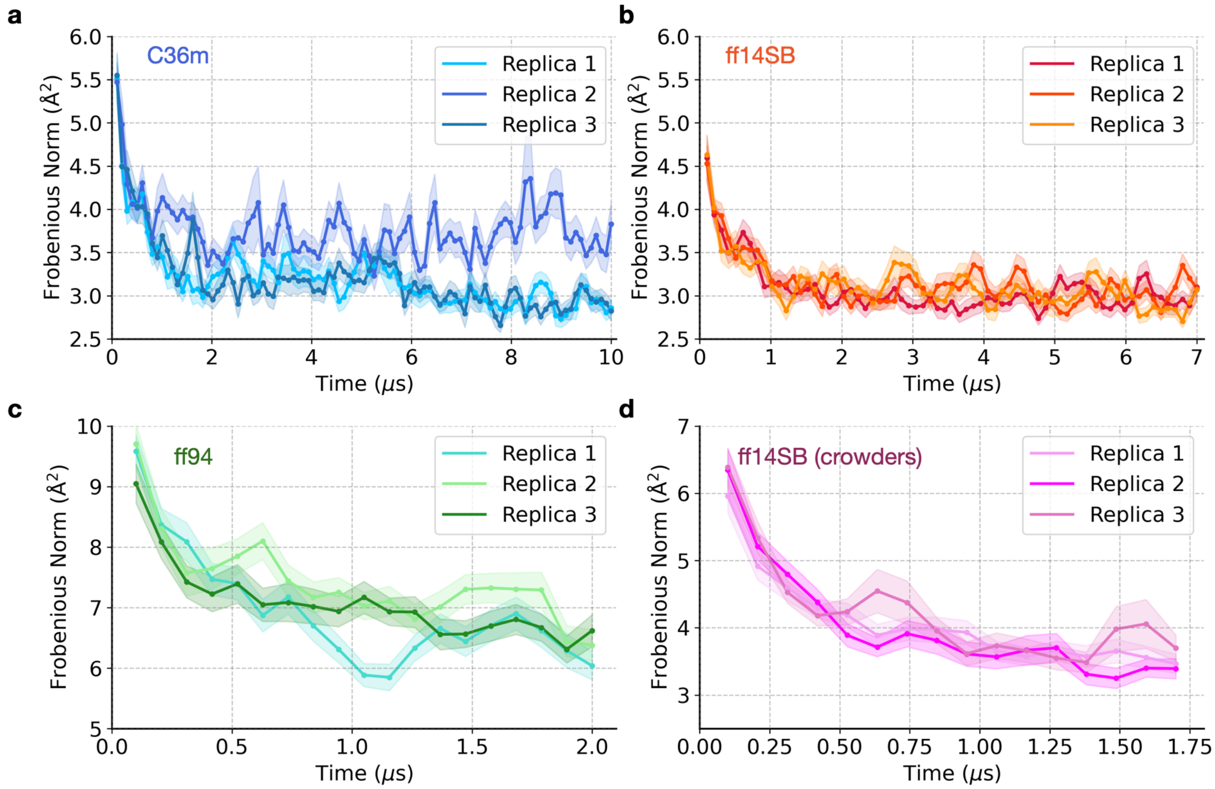

**Supplementary Figure** 3: **Convergence analysis of atomic covariance matrices**. Covariance matrices $\text{cov}_{\beta\gamma}$ capturing the relationships between pairs of $C\alpha$ atom coordinates $\beta, \gamma$ were computed within each 100 ns time window. Convergence was assessed by examining the distance between covariance matrices taken at time $t$ and $t + 100$ ns. This distance, measured by the Frobenius norm $(||\text{cov}_{\beta\gamma}(t+100) - \text{cov}_{\beta\gamma}(t)||)$, was averaged over $n = 108$ chains and depicted as a function of time $t$ (in $\mu$s) for supercell simulations; in the simplified crystal environment using force fields (**a**) C36m, (**b**) ff14SB, and (**c**) ff94, as well as (**d**) ff14SB with crowders. The standard error ($n = 108$) for each replica is represented by a shaded envelope. Note that the simulations with (**c**) ff94, and (**d**) ff14SB with crowders have not reached equilibrium. For further details on the calculation of atomic covariance matrices, refer to SI section 3.4.

#### 2.3 Other crystal structures of the second PDZ domain of LNX2

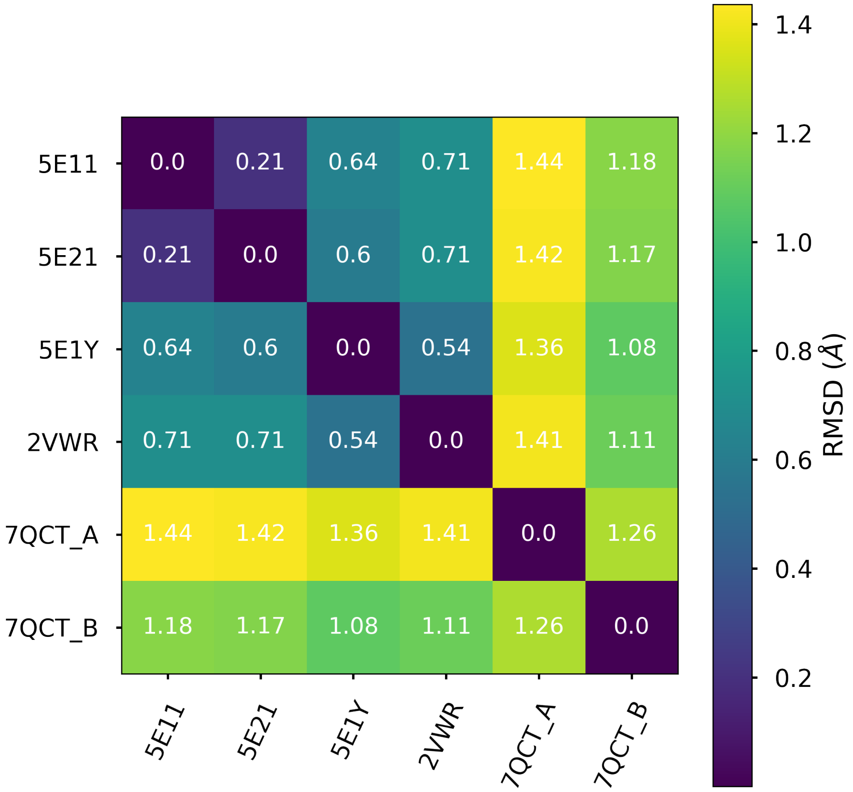

**Supplementary Figure** 4: **Deviation between crystal structures of the second PDZ domain of LNX2.** The pairwise RMSD between all available crystal structures of the second PDZ domain of the human LNX2 protein is shown. RMSD was computed using heavy atoms in the residues common to all crystal structures (residues 336 to 424, excluding 338, in the numeration of PDB ID: 5E11). The CHARMM-GUI web server[25] was used to add missing atoms.

#### 2.4 B-factors for ff94 and ff14SB+crowders simulations

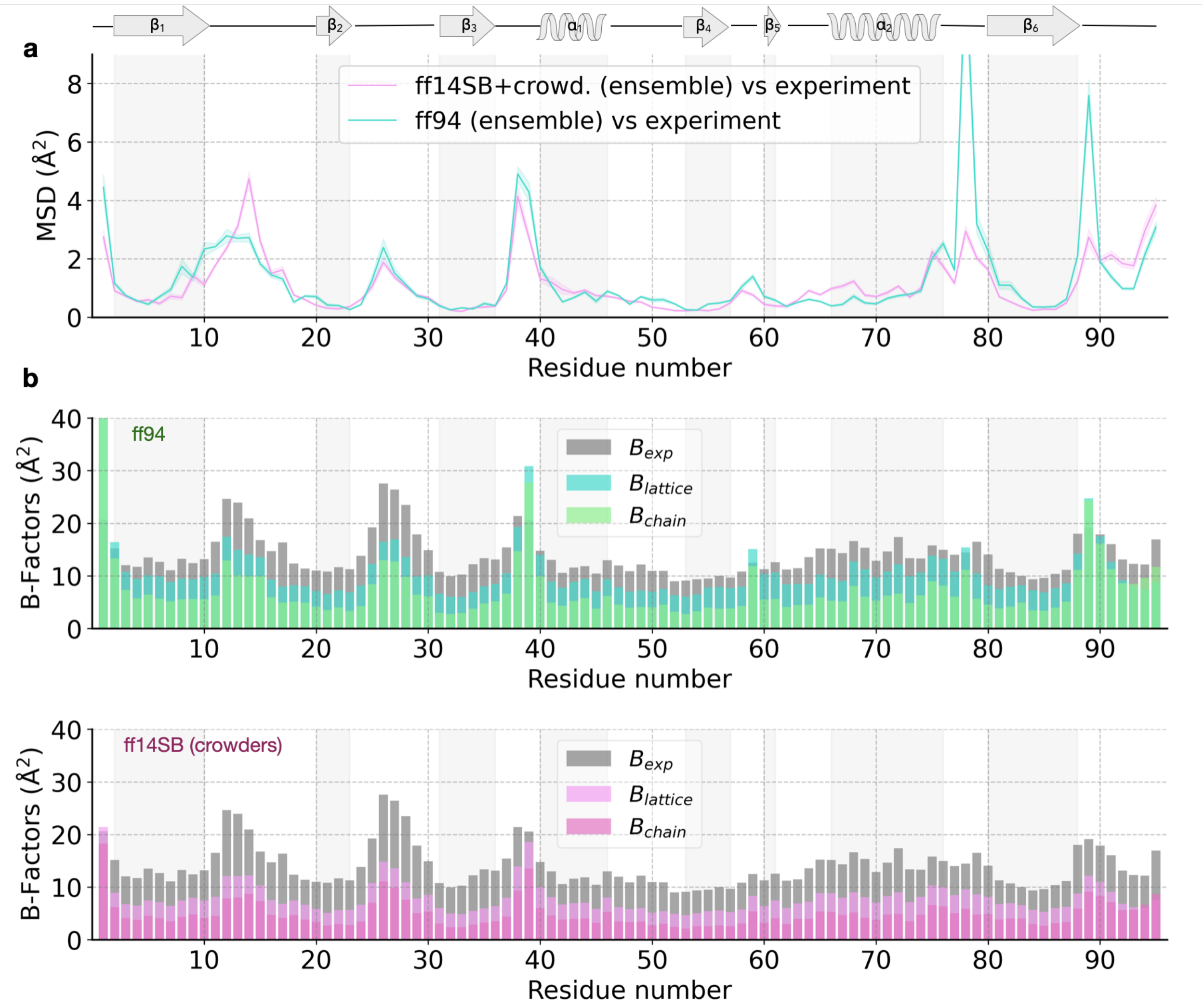

**Supplementary Figure** 5: **Comparison to the crystal structure and B-factors for ff94 and ff14SB with crowders simulations. (a)** MSD of C$\alpha$ positions relative to the crystal structure (PDB ID: 5E11), with the shaded envelope representing the mean $\pm$ standard deviation ($n$ = 324 chains = 3 replicas $\times$ 108 copies). The ff94 simulations show a high deviation in the $\alpha_2$-$\beta_6$ region and the C-terminal tail. **(b)** Comparison between simulation ($B_{\text{lattice}}$ and $B_{\text{chain}}$) and experimental ($B_{\text{exp}}$) B-factors computed for C$\alpha$ atoms. Note that bars are overlaid, not stacked. The correlation with the experimental B-factors is high for the ff14SB+crowders simulations (Pearson $r$ = 0.84 and 0.80 for $B_{\text{lattice}}$ and $B_{\text{chain}}$, respectively), while the correlation is lower for the ff94 simulations (Pearson $r$ = 0.59 and 0.55 for $B_{\text{lattice}}$ and $B_{\text{chain}}$, respectively).

#### 2.5 Liquid-like motions in the crystal

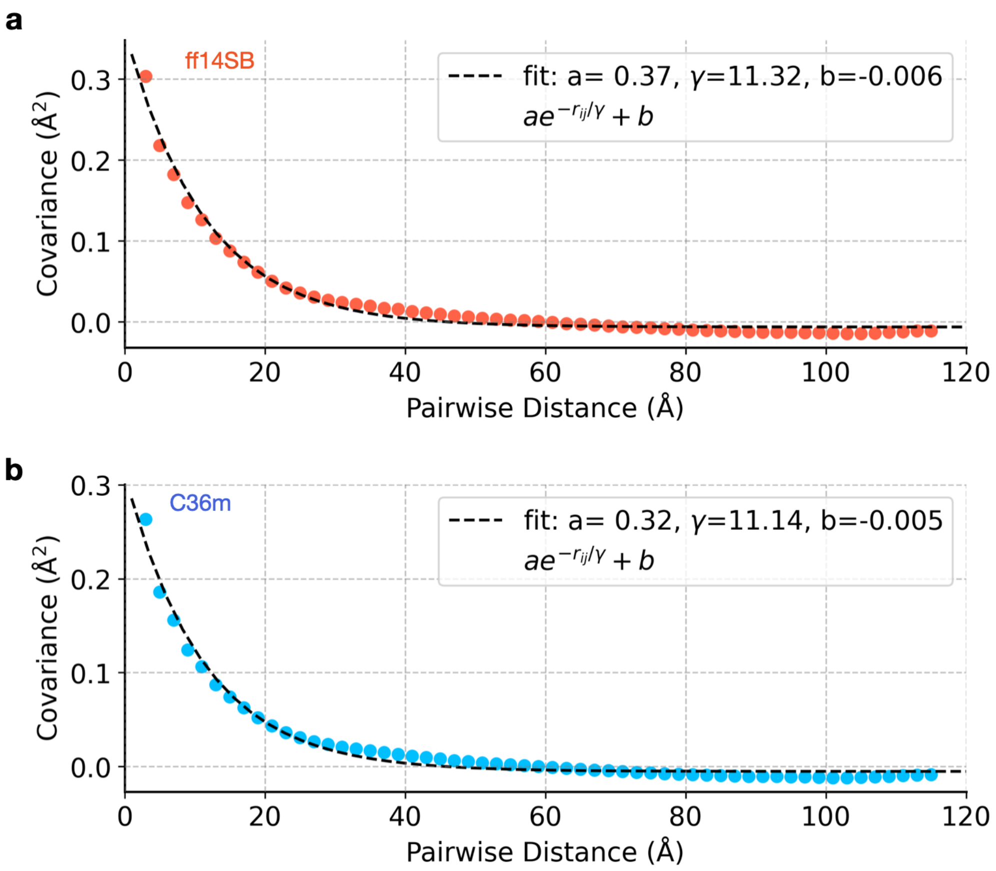

**Supplementary Figure 6: Dependence of atomic covariance on the inter-atomic distance.** The covariance and distance matrices were computed for all C$\alpha$ atoms in a supercell. To plot covariance vs. distance, the elements of the distance matrix were distributed into 60 equal bins in a range from 0 to 120 Å and covariance matrix elements for pairs of atoms within each bin were averaged. The vertical error bars are smaller than the size of the markers. The data was fit to an exponential function $\mathrm{cov}_{ij}(r_{ij}) = ae^{-r_{ij}/\gamma} + b$. **(a)** For the ff14SB force field, the values of parameters are $a = 0.37 \pm 0.02$ Å$^2$, $\gamma = 11.3 \pm 0.7$ Å and $b = -0.006 \pm 0.003$ Å$^2$. **(b)** For the C36m force field, the values of parameters are $a = 0.32 \pm 0.02$ Å$^2$, $\gamma = 11.1 \pm 0.8$ Å and $b = -0.005 \pm 0.002$ Å$^2$. The error estimate provided for each fitting parameter is the 95% confidence interval. For further details regarding the method of computing atomic covariance, refer to SI section 3.4.

##### 2.6 Crystal lattice disorder

In order to determine how much protein chains deviate from their ideal positions in the crystal lattice, we quantified the crystal lattice disorder. Inverse crystallographic transformations (described below) were applied to the positions of each of the 108 protein chains in the supercell simulations. In the ideal case of an undistorted lattice, all points would remain at the origin. Due to thermal motion and melting symmetry (described in detail in SI section 2.7), the positions of the center of mass of individual chains scatter around their undistorted positions, which would be at the origin once the inverse crystallographic transformations are applied. The amplitude of these deviations from the origin indicates the level of disorder in the crystal lattice. The deviation at each frame was computed as $\left(\frac{1}{n}\sum_{i=1}^{n} x_i^2(t) + y_i^2(t) + z_i^2(t)\right)^{\frac{1}{2}}$, where $x_i, y_i, z_i$ are center-of-mass positions of the $n = 108$ individual protein chains in the inversely transformed lattice. The average value of this deviation from the origin over the last 1 $\mu$s of simulation for each replica is provided in Supplementary Table 2.

**Inverse crystallographic transformations**. For the crystal lattice model, a $3 \times 3 \times 3$ supercell was constructed from the experimental PDB structure (PDB ID: 5E11). The coordinates of each chain were obtained by linear transformations applied to the original PDB coordinates. These operations (rotations and translations) are defined by the crystal symmetry group ($C121$) and the unit cell dimensions ($a = 65.30$ Å, $b = 39.45$ Å, $c = 39.01$ Å and $\alpha = \gamma = 90°$, $\beta = 117.54°$). The inverse crystal transformations are found using $\hat{A}_{\text{inv}}(t) = \hat{A}(t)^{-1}$, where transformations $\hat{A}(t)$ depend on time as the unit cell axes change due to the pressure coupling algorithm (Supplementary Figs. 7-9), which is described in detail in SI section 2.7.

To visualize lattice dynamics, the center of mass positions of the transformed chains were plotted for each simulation frame. The instantaneous centers-of-mass of all 108 protein chains were projected onto the **ac** crystallographic plane. Supplementary Movies 5 and 6 show visualizations of lattice dynamics in the simulations of the PDZ domain supercell using

**Supplementary Table 2: Quantifying lattice disorder in the supercell.** For each simulation of the supercell, the time averaged deviation from the undistorted position (computed as described above) is provided. Replicas that have not reached equilibrium are indicated with ($\star$) and are not used to compute the average lattice deviation reported in the main text (1.61 Å for ff14SB and 1.82 Å for C36m).

| Force Field | Replica | Time-averaged deviation from undistorted position in the last $\mu$s (Å) |
|---|---|---|
| ff14SB | 1 | 1.62 |
| | 2 | 1.59 |
| | 3 | 1.62 |
| C36m | 1 | 1.86 |
| | $2^{\star}$ | 1.27 |
| | 3 | 1.76 |
| ff94 | $1^{\star}$ | 1.63 |
| | $2^{\star}$ | 1.87 |
| | $3^{\star}$ | 1.67 |
| ff14SB + crowd. | $1^{\star}$ | 1.76 |
| | $2^{\star}$ | 1.84 |
| | $3^{\star}$ | 1.77 |

the ff14SB and C36m force fields. In these animations, the two clusters of points (visible in the NPT simulation, i.e. after $t = 20$ ns of pre-production) represent chains in two different orientations that are shifted relative to each other (exhibiting melting symmetry, as described in SI section 2.7).

#### 2.7 Melting crystal symmetry

Melting crystal symmetry is observed in the crystal simulations when simulation box vectors (i.e. crystallographic axes) change due to the effects of the anisotropic pressure coupling algorithm in the NPT ensemble. To quantify this effect, we analyzed various properties of the protein crystal as a function of simulation time and simulation protocol (Supplementary Figs. 7 and 8). We note that all simulation replicas were initialized from the same coordinates but with different starting velocities.

The most significant conformational change happens in the pre-production, during the NVT run, after the position restraints are lifted ($t = 0$ to 10 ns, green area in Supplementary Figs. 7 and 8). A rapid increase of the RMSD occurs even when supercell dimensions are still constant (from 0.4 to 1.1 Å). The radius of gyration increases as well, but the change in the protein structure occurs anisotropically; the protein contracts along axis **b** and extends along axis **c** (Supplementary Fig. 7). This is the case for simulations using both force fields (ff14SB and C36m).

After switching to the isotropic Berendsen NPT ensemble ($t = 10$ to 20 ns, pink area in Supplementary Figs. 7 and 8), the system volume stays similar to the experimental volume, never deviating more than 0.25% (Supplementary Fig. 8). To maintain constant pressure, a significant rescaling of unit cell vectors occurs ($a_X$, $b_Y$, $c_Z$ in Supplementary Fig. 8), which is typical for MD simulations of protein crystals in the NPT ensemble.[26] Such rescaling of box vectors continues the trend of anisotropic deformation of the protein ($Rg_Y$, $Rg_Z$ in Supplementary Fig. 7), but the changes are less significant than those observed in the first 10 ns. Importantly, this causes a relative center of mass drift of individual chains, which we refer to as "melting symmetry" ($\mathrm{COM}_X$, $\mathrm{COM}_Y$, $\mathrm{COM}_Z$ in Supplementary Figs. 8 and 9). For instance, in simulations with ff14SB, we observed a center of mass distance between adjacent chains in nearest-neighbour unit cells to increase by 0.5 Å along **a** and by 0.8 Å along **c**, as well as a decrease by 1.2 Å along **b**. Similar melting symmetry effects were observed in the C36m simulations.

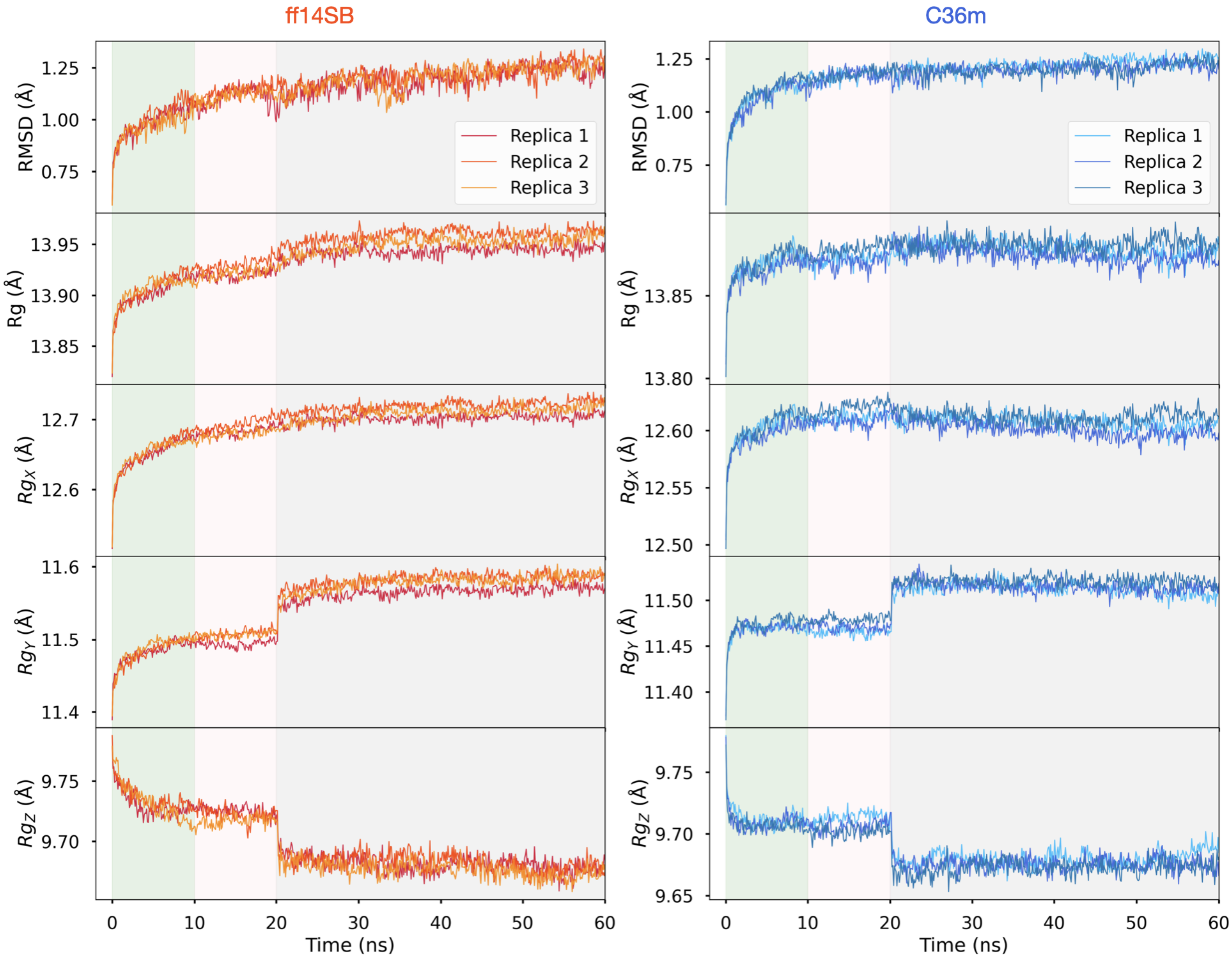

**Supplementary Figure 7: Structural observables (RMSD and $Rg$) during pre-production runs.** The first 10 ns is the NVT equlibration (green region), followed by 10 ns of isotropic Berendsen NPT (pink region), followed by 40 ns of anisotropic Parinello-Rahman NPT (gray region). The average RMSD from the crystal structure (PDB ID: 5E11) for heavy atoms, average radius of gyration $Rg$, as well as radii of gyration ($Rg_X$, $Rg_Y$, $Rg_Z$) along axes $X, Y, Z$ are plotted vs. time. The average values are computed for $n = 108$ chains in each replica.

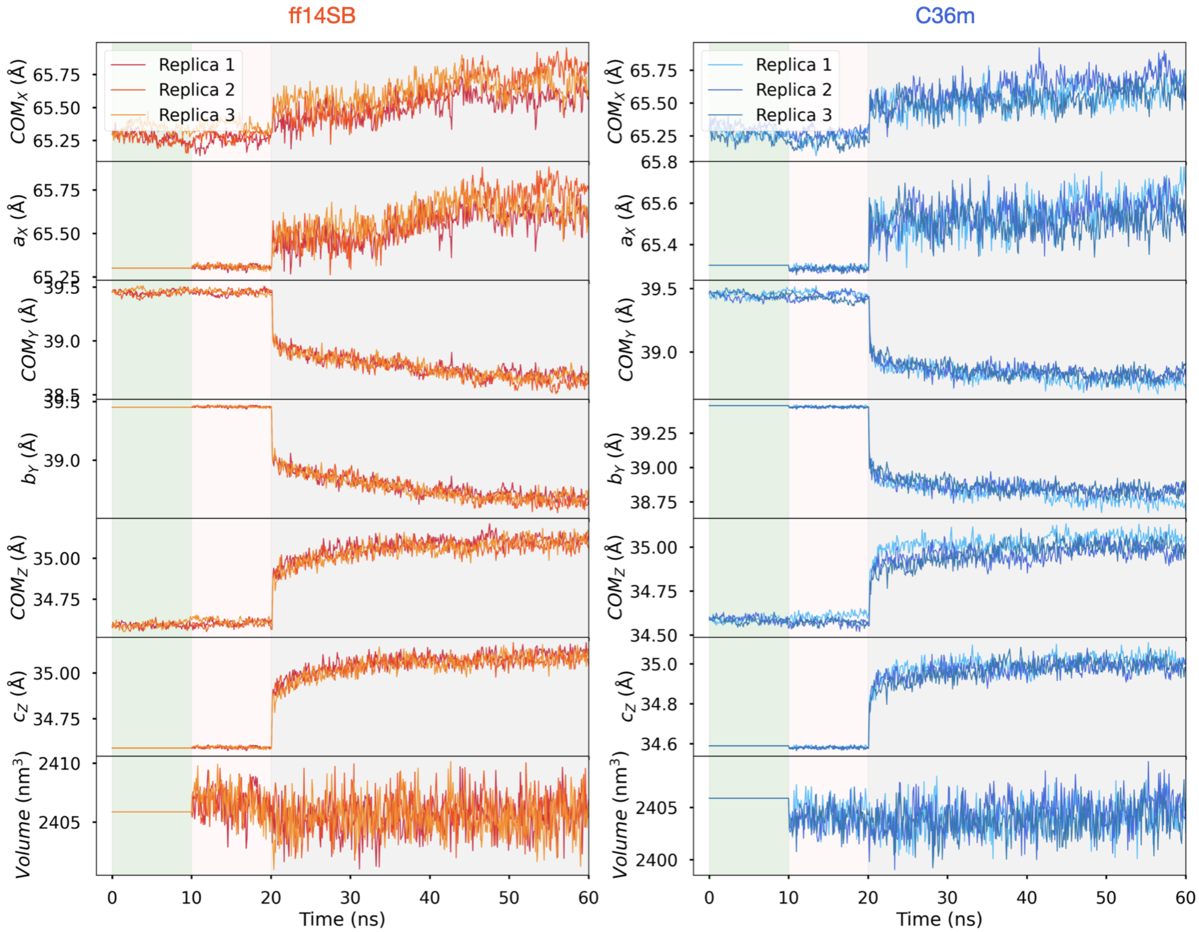

**Supplementary Figure** 8: **Unit cell axes and distance between protein chains in pre-production.** The simulation protocol regions (NVT, isotropic NPT Berendsen and anisotropic NPT Parinello-Rahman) are coloured as in Supplementary Fig. 7. $\mathrm{COM}_X$, $\mathrm{COM}_Y$, $\mathrm{COM}_Z$ are the $X, Y, Z$ components of the average distance between the protein chains (center of mass) in the nearest-neighbour unit cells along crystallographic vectors **a**, **b**, **c**, respectively. $a_X$, $b_Y$, $c_Z$ are the $X, Y, Z$ components of the crystallographic vectors, respectively. System volume is plotted in the last row.

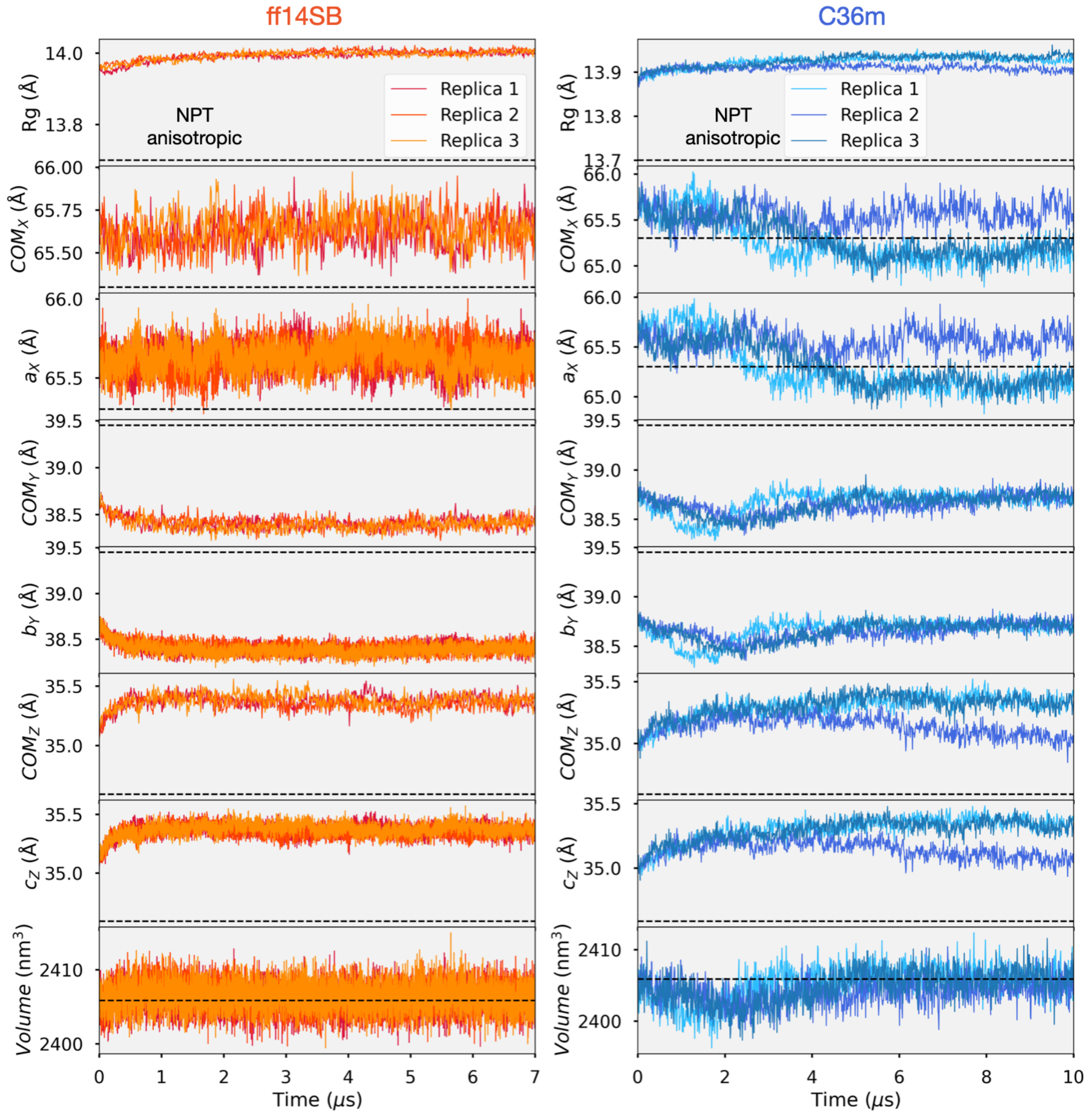

**Supplementary Figure 9: Radius of gyration, unit cell axes and distance between protein chains during the production simulations.** $\mathrm{COM}_X$, $\mathrm{COM}_Y$, $\mathrm{COM}_Z$ are the $X, Y, Z$ components of the average distance between the chains (center of mass) in the nearest-neighbour unit cells along crystallographic vectors **a**, **b**, **c**, respectively. $a_X$, $b_Y$, $c_Z$ are the $X, Y, Z$ components of the crystallographic vectors, respectively. System volume is plotted in the last row. Dashed lines represent experimental values.

#### 2.8 Inter-protein contacts

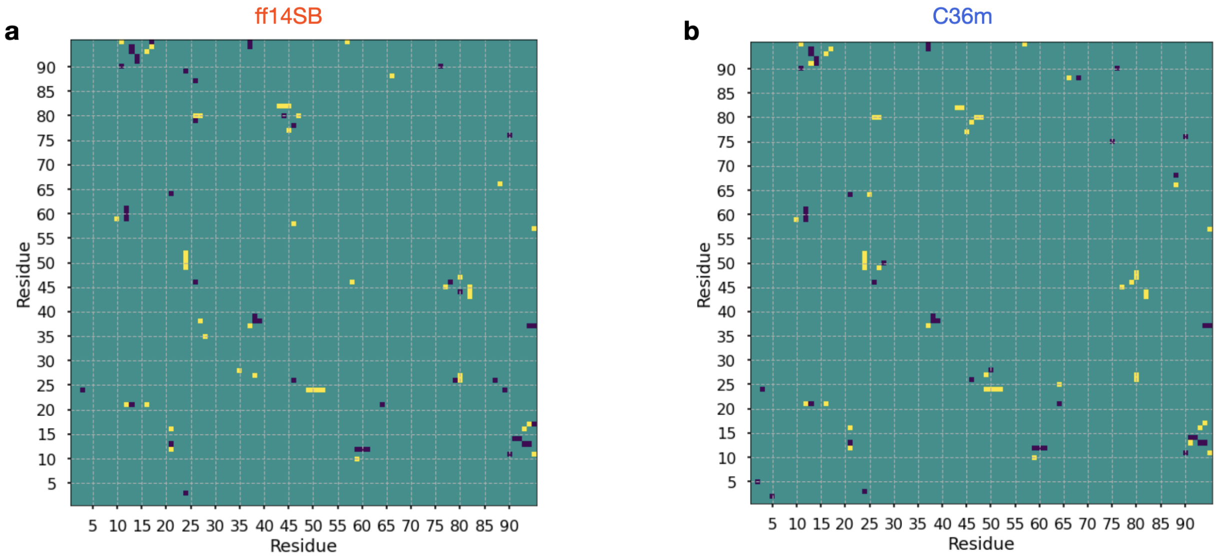

**Supplementary Figure** 10: **Inter-protein contact maps — difference between simulations and experiment.** The difference contact maps for **(a)** ff14SB and **(b)** C36m are shown. Yellow elements represent the new contacts formed in simulation, blue elements represent the contacts lost in the simulation, and green indicates no change compared to the crystal structure (PDB ID: 5E11). Only the contacts with a propensity > 0.5 are shown. In the crystal structure, 69 out of 95 protein residues are involved in inter-molecular contacts. In simulations, there are 23 broken and 23 newly formed inter-protein contacts for ff14SB, and there are 21 broken and 24 newly formed for C36m.

**Supplementary Table 3: Segments involved in lost and newly formed inter-protein contacts in both ff14SB and C36m.** The segments that experienced contact interaction changes compared to the crystal structure (PDB ID: 5E11) were associated with significant conformational changes; these regions include the C-terminal ligand motif, $\beta_1$-$\beta_2$, and $\beta_3$-$\alpha_1$, as well as the most flexible regions, such as $\beta_2$-$\beta_3$. Other changes in the crystal contacts occurred in less flexible segments due to side chain rearrangements. One such instance involves the $\beta_6$ strand (residues 80-82), which forms new crystal contacts to the $\alpha_1$ helix (residues 43-45), which is known to be allosterically coupled to the ligand-binding site.[27–30] Thus, all changes to the inter-protein interactions were either related to the deviations in the protein structure or involved functionally relevant regions of the PDZ domain.

| # | lost/formed | Segment 1 | Segment 2 | Residues involved |
|---|---|---|---|---|
| 1 | lost | $\beta_1$-$\beta_2$ loop | C-term. | 12-14 — 93-94 |
| 2 | formed | $\beta_1$-$\beta_2$ loop | C-term. | 16-17 — 93-94 |
| 3 | lost | $\beta_1$-$\beta_2$ loop | $\beta_5$ strand | 12-14 — 59-61 |
| 4 | formed | $\beta_1$-$\beta_2$ loop | $\beta_5$ strand | 10 — 59-61 |
| 5 | formed | $\beta_6$ strand | $\beta_2$-$\beta_3$ loop | 80-82 — 27-28 |
| 6 | formed | $\beta_6$ strand | $\alpha_1$ helix | 80-82 — 43-45 |
| 7 | formed | $\beta_2$-$\beta_3$ loop | $\alpha_1$-$\beta_4$ loop | 24 — 49-52 |
| 8 | lost | $\beta_3$-$\alpha_1$ loop | $\beta_3$-$\alpha_1$ loop | 38-39 — 38-39 |
| 9 | formed | $\beta_3$-$\alpha_1$ loop | $\beta_3$-$\alpha_1$ loop | 37 — 37 |

#### 2.9 Analysis of crystallographic water sites

**Supplementary Table 4: The percentage of water sites classified as preserved, bulk-like and obstructed in crystal simulations.** The LAWS algorithm[31] was used to carry out this analysis. The replicas that have not reached equilibrium are indicated with (⋆) and are not used to compute the average CWS recall reported in the main text (71% for ff14SB and 71% for C36m).

| Force Field | Replica | Preserved CWS (%) | Bulk-like WS (%) | Obstructed WS (%) |
|---|---|---|---|---|
| ff14SB | 1 | 71 | 19 | 10 |
| | 2 | 72 | 18 | 10 |
| | 3 | 69 | 21 | 10 |
| C36m | 1 | 72 | 15 | 13 |
| | 2⋆ | 76 | 11 | 14 |
| | 3 | 70 | 22 | 8 |
| ff94 | 1⋆ | 50 | 32 | 18 |
| | 2⋆ | 53 | 35 | 12 |
| | 3⋆ | 59 | 30 | 11 |
| ff14SB + crowd. | 1⋆ | 70 | 21 | 9 |
| | 2⋆ | 70 | 21 | 9 |
| | 3⋆ | 71 | 22 | 6 |

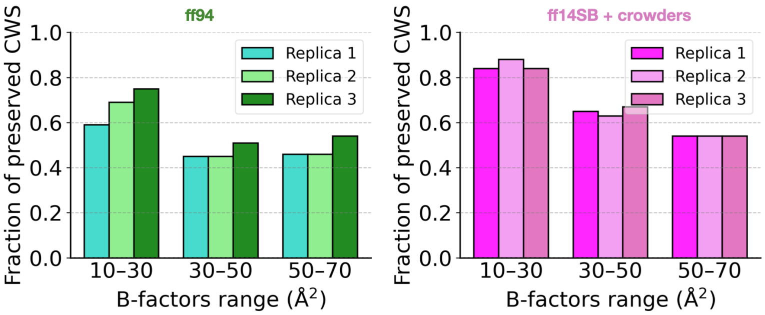

**Supplementary Figure 11: The fraction of preserved crystallographic water sites in the ff94 and ff14SB + crowders simulations grouped by experimental B-factor.** All 94 crystallographic water sites are binned into 3 groups according to their experimenal B-factor: 10–30 Å$^2$ ($n = 32$), 30–50 Å$^2$ ($n = 49$), and 50–70 Å$^2$ ($n = 13$). Distributions for ff14SB and C36m are provided in the main text (Fig. 2e).

**Supplementary Table 5: Protein segments associated with lost crystallographic waters.** The LAWS algorithm[31] was used to assess whether or not each CWS in the crystal structure was preserved in the simulation. Twelve CWS were identified as not preserved in both ff14SB and C36m simulations. Of these 12 CWS, 4 are coordinated by the residues of a single chain (interior), while 8 are coordinated by segments of multiple chains (interface). These CWS are labeled (first column) with numbers according to the numeration of water sites in the crystal structure (PDB ID: 5E11). The ligand binding site of the PDZ domain is a pocket between the $\alpha_2$-helix and the $\beta_2$-strand.

| # | Type | Location |
|---|---|---|
| 44 | interior | within the $\beta_3$-$\alpha_1$ loop |
| 49 | interior | between $\alpha_2$-$\beta_6$ and $\beta_1$-$\beta_2$ loops |
| 74 | interior | between $\alpha_2$-$\beta_6$ and $\beta_1$-$\beta_2$ loops |
| 91 | interior | between side chains of Gln72 and Ile73 ($\alpha_2$) |
| 1 | interface | between $\beta_2$-$\beta_3$ loop and $\alpha_1$ helix and $\alpha_2$-$\beta_6$ loop |
| 5 | interface | between $\beta_1$-$\beta_2$ loop and $\beta_3$-$\alpha_1$ loop |
| 7 | interface | between $\alpha_2$-helix and $\beta_2$-$\beta_3$ loop |
| 9 | interface | between $\beta_1$-$\beta_2$ loop and C-term. (binding site) |
| 17 | interface | between $\alpha_2$-helix and C-term. (binding site) |
| 75 | interface | between $\alpha_1$-helix and $\beta_4$-$\beta_5$ turn |
| 77 | interface | between $\beta_1$-strand and $\beta_6$-strand and $\alpha_2$-helix |
| 93 | interface | between $\beta_2$-$\beta_3$ loop and $\alpha_1$-helix |

#### 2.10 Principal component analysis — alternative featurizations

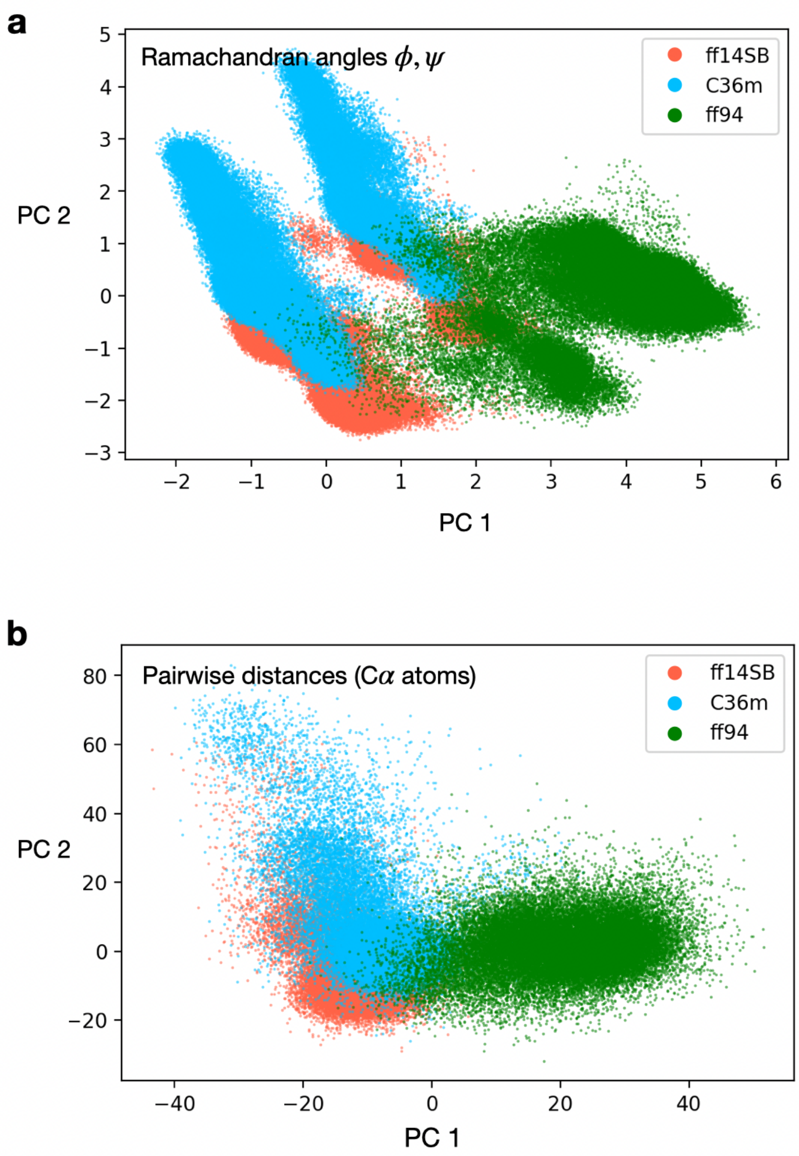

**Supplementary Figure** 12: **Principal component analysis of the crystalline simulations using alternative featurizations.** The two-dimensional PCA projection of the conformational ensembles obtained with various force fields, where **(a)** Ramachandran angles (sin/cos transformed),[32] and **(b)** pairwise distances for C$\alpha$ atoms are used as features.

#### 2.11 Principal component analysis — importance of individual residues

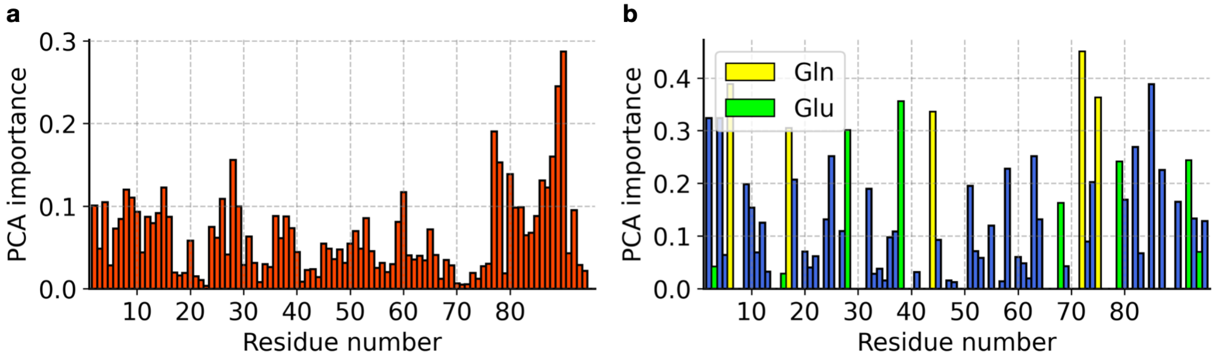

**Supplementary Figure** 13: **Importance of individual residues in the first principal component.** The PCA importance is shown for principal component 1 (PC 1) in the PCA reported in Fig. 3a. PCA importance is computed as a sum of magnitudes of coefficients in PC 1 corresponding to the specific residues for **(a)** $\phi$, $\psi$ Ramachandran angles, **(b)** $\chi_1$, $\chi_2$ Janin angles, in the feature vector. All five glutamine residues (yellow) are in the top 10 residues with highest importance. Several glutamic acid residues (green) also have high PCA importance.

#### 2.12 Analysis of the ff14SB and C36m ensembles using LDA and PCA

The analysis of the conformational space of the PDZ domain in the crystalline state shows that the conformational space is force field dependent, i.e., the space is clustered into three regions corresponding to the three force fields (ff14SB, C36m, and ff94) used for the simulations (Fig. 3a in the main text). Here, we aim to understand the cause of the differences between ff14SB and C36m specifically, as these two force fields are the most recent and widely used. To highlight the specific differences in the conformational space sampled between ff14SB and C36m, we excluded the ff94 ensemble from the analysis. Principal component analysis focuses on maximizing the variance in the data by transforming it into a new set of uncorrelated variables, the principal components. While these components may capture the most significant sources of variation, they are not necessarily oriented to optimize class discrimination. Conversely, linear discriminant analysis (LDA) is explicitly designed for supervised classification tasks.[33] It seeks to find linear combinations of features that maximize the separation between different classes. LDA takes into account class information, aiming to reduce within-class variability and enhance between-class variability.

Here, we carry out both types of analysis using both Ramachandran and Janin normalized (sin/cos) angles as features (Supplementary Fig. 14), which is the same featurization used in the PCA shown in Fig. 3a of the main text. Protein conformations were sampled every 10 ns from the ff14SB and C36m crystal simulation trajectories (Table 1 of the main text), resulting in 528,876 protein conformations in total. Class information (i.e. force field) was used for LDA. The *sklearn 1.2.0* Python package[34] was used to carry out PCA and LDA.

**PCA**. The results of pairwise PCA are consistent with the analysis where the ff94 ensemble was included (Fig. 3a); the force field regions are split along the PC axis 1 (Supplementary Fig. 14a). Analyzing the importance of residues in PC 1, we find that the glutamine (resid 6, 17, 44, 72, 75), glutamic acid (resid 28, 38, 92), methionine (resid 2), and isoleucine (resid 85) residues are in the list of the top 10 important residues. All five (out of 5) glutamines

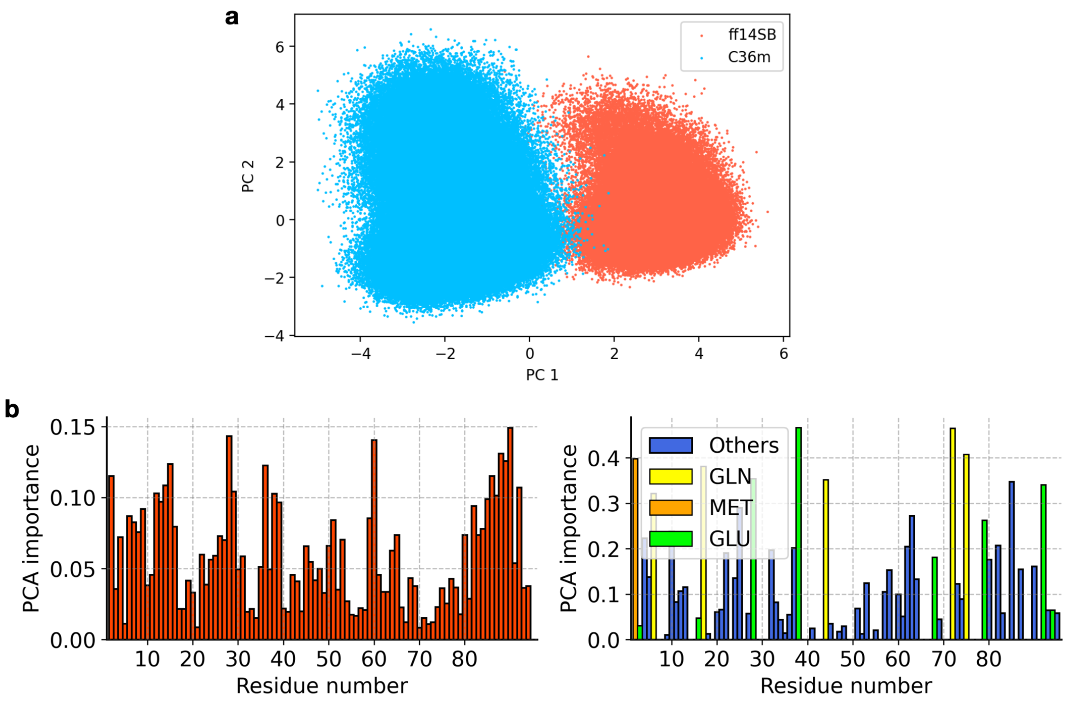

**Supplementary Figure** 14: **PCA of the ff14SB and C36m ensembles**. **(a)** Projection of the simulated conformations on the PCA space; the conformational ensemble is separated into two force field basins. The same feature vectors as in the PCA shown in Fig. 3a were used. **(b)** The PCA importance is shown for PC 1. PCA importance is computed as a sum of magnitudes of coefficients in PC 1 corresponding to the specific residues for $\phi$, $\psi$ Ramachandran angles (left) and $\chi_1$, $\chi_2$ Janin angles (right) in the feature vector. As in Supplementary Fig. 13, all five glutamine residues (yellow) are in the top 10 important residues.

are in the top 10 residues, while three (out of 8) glutamic acid residues, a single isoleucine (out of 8), and a single methionine (out of 1) are in this list (Supplementary Fig. 14b).

**LDA**. The results of LDA are consistent with PCA; the force field regions are linearly separable along the LDA axis (Supplementary Fig. 15a). Analyzing the importance of residues, we find that the glutamic acid (resid 28, 38, 79, 68), glutamine (resid 72, 75), methionine (resid 2), lysine (resid 21) and histidine (resid 10) residues are in the list of the top 10 important residues for the side chain conformation (sum of $\chi_1$ and $\chi_2$ coefficients). Specifically, glutamic acid (6 out 8) and glutamine (4 out of 5) residues have the highest contribution to the separation of the force fields in terms of $\chi_2$ dihedrals (Supplementary Fig. 15b, right).

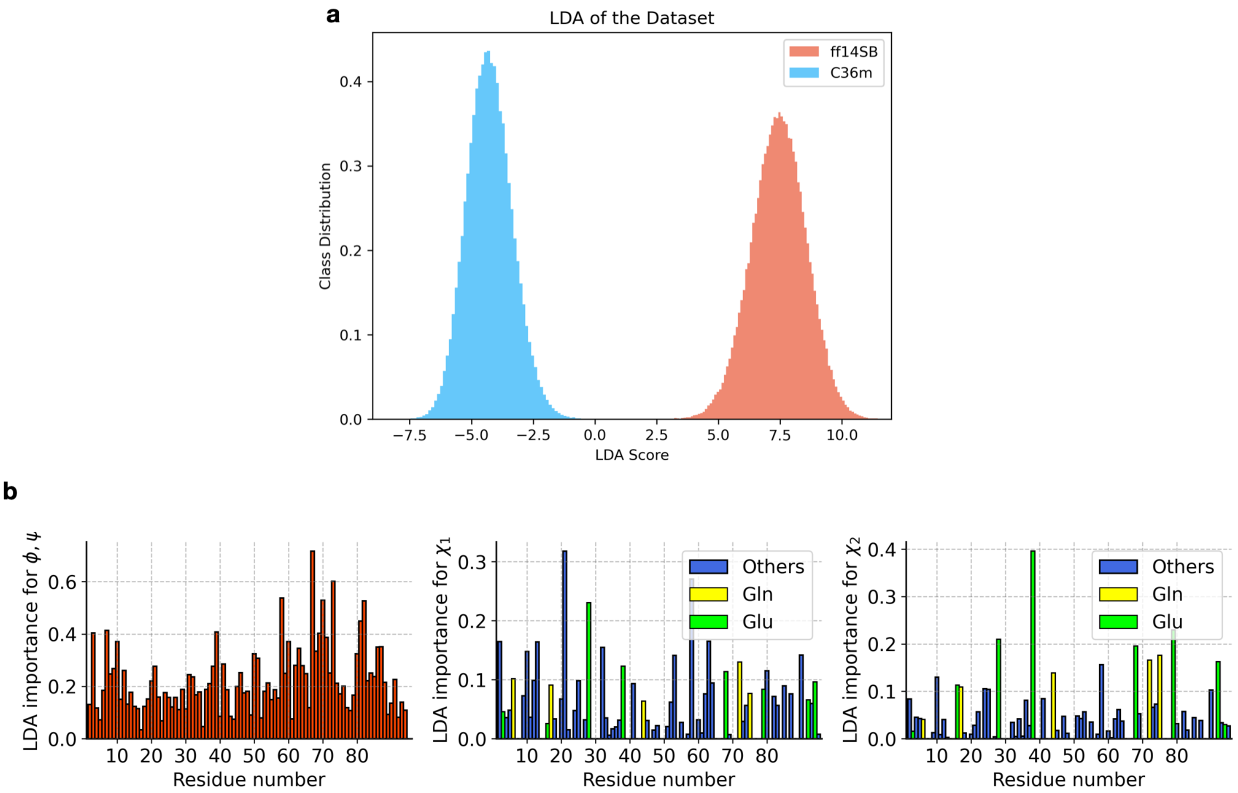

**Supplementary Figure** 15: **LDA of the ff14SB and C36m ensembles**. **(a)** The distribution of the ff14SB and C36m ensembles along the single LDA axis; the conformational ensemble is separated into two force field basins. The same featurization as the PCA shown in Fig. 3a was used. **(b)** The LDA importance is computed as a sum of magnitudes of coefficients of the first discriminant corresponding to the specific residues for $\phi$,$\psi$ (left), $\chi_1$ (center), and $\chi_2$ angles (right) in the feature vector.

We, therefore, conclude that side chains of glutamine and glutamic acid residues are likely the primary cause of the difference between ff14SB and C36m. From this analysis, we do not have sufficient statistics to generalize conclusions about methionine, lysine, histidine and isoleucine.

##### 2.13 Side chain rotamers of residues with low PCA importance

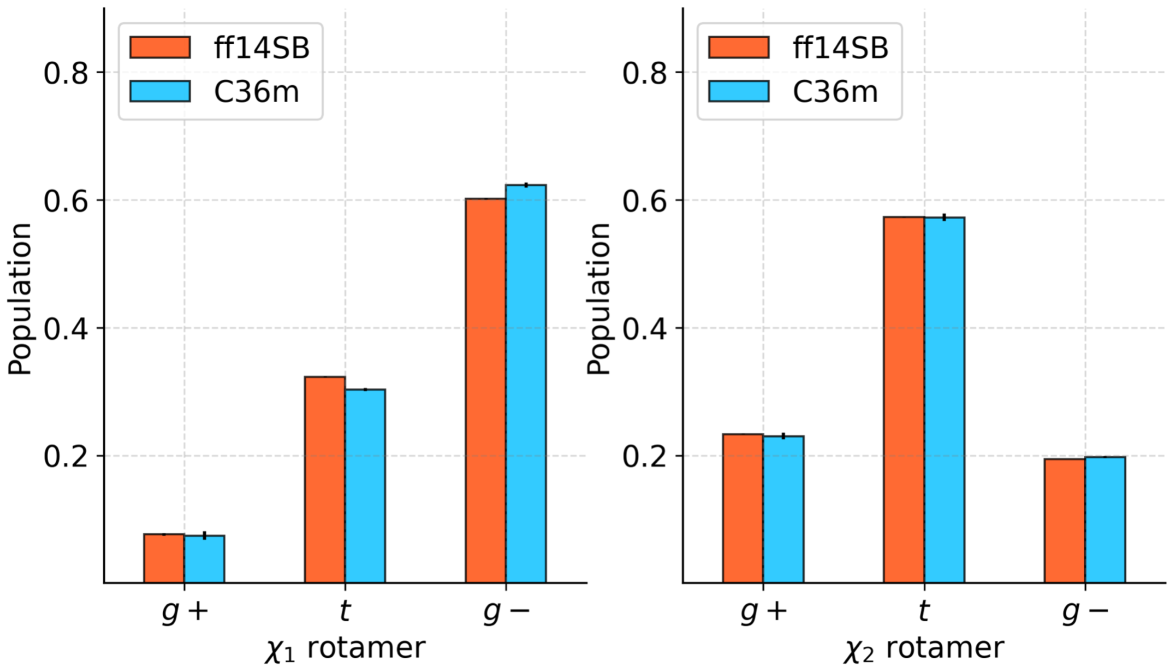

**Supplementary Figure** 16: **Combined distribution of side chain rotamers** ($\chi_1$, $\chi_2$) **in residues with low PCA importance**. This analysis includes the bottom 50 out of 60 residues in side chain PCA importance (shown in Supplementary Fig. 13b). The distributions are sampled from the ff14SB and C36m crystal simulations and standard errors are calculated for $n = 3$ replicas.

#### 2.14 Glutamine and glutamic acid side chain rotamers in other simulation systems

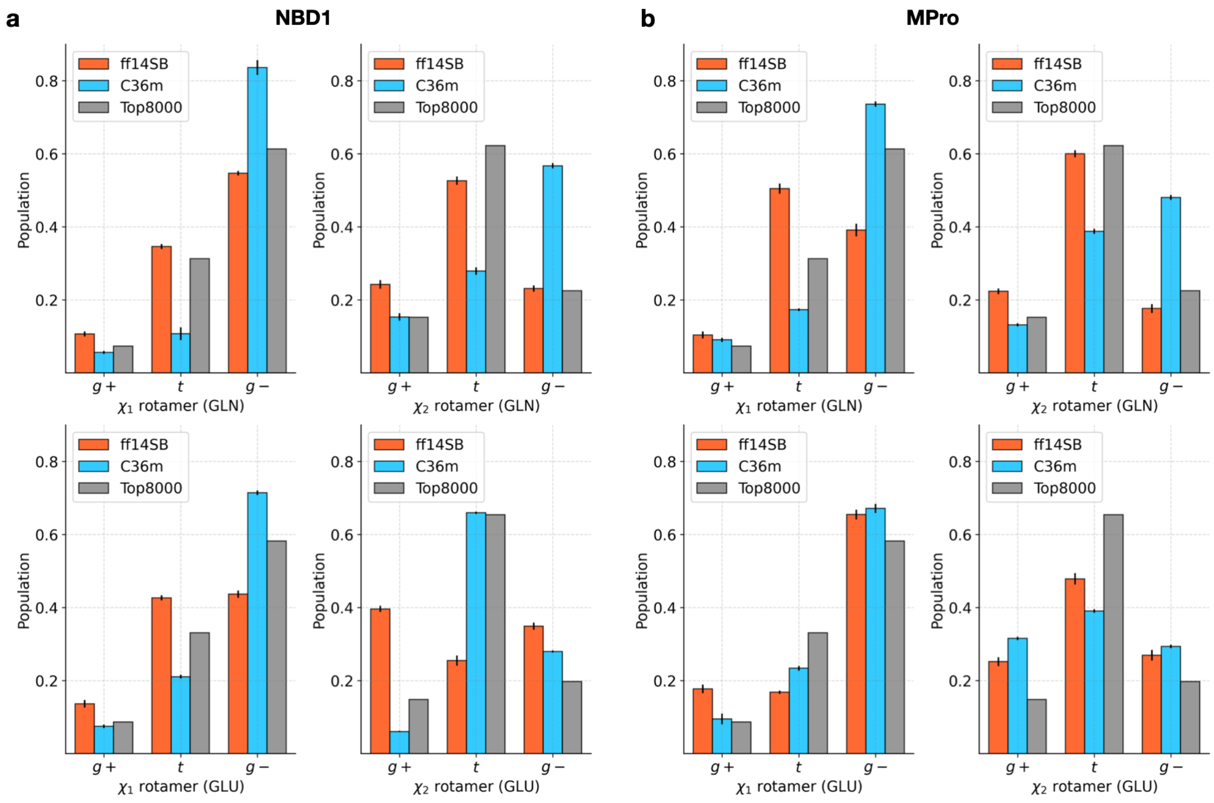

**Supplementary Figure** 17: **Distribution of $\chi_1$, $\chi_2$ rotamers for glutamine and glutamic acid in other simulation systems.** Distributions are shown for two additional protein systems, NBD1 and $M^{pro}$, simulated using both the ff14SB and C36m force fields. The distributions sampled from the Top8000 dataset are shown in grey. Standard errors are computed for $n$ simulation replicas in each system (refer to the simulation protocol described above). In NBD1, there are 15 glutamines and 10 glutamic acid residues. In $M^{pro}$, there are 28 glutamines and 18 glutamic acid residues.

We carried out the analysis of side chain rotamers in two other simulation systems, a nucleotide binding domain of a sulfonylurea receptor (NBD1 of the splice isoform of human SUR2 with exon 17 deleted,[35] 238 residues) and the main protease of SARS-CoV[36] ($M^{pro}$, 612 residues). NBD1 is the site of MgATP binding in the human sulfonylurea receptor, which regulates the opening and closing of ATP sensitive potassium channels ($K_{ATP}$ channels).[35] The main protease ($M^{pro}$) is one of two essential proteases within the SARS-CoV genome that cleaves polyproteins expressed during viral replication.[36] Similarly to the PDZ domain system, we computed the distributions of $\chi_1$ and $\chi_2$ rotamers for all glutamine and glutamic acid residues in the $M^{pro}$ and NBD1 trajectories using the ff14SB and C36m force fields and compared these to the distributions from the Top8000 dataset[37] (Supplementary Fig. 17). These results demonstrate that in these two systems, the distribution of $\chi_1$ and $\chi_2$ rotamers for glutamine residues differ between force fields (as observed for the PDZ domain simulations), while the differences for glutamic acid residues are less pronounced.

**Simulation protocol.** The simulation setup for these two systems is similar to the one used for the PDZ domain in solution. The differences are as follows. For the $M^{pro}$ and NBD1 systems, 0.15 M concentration of NaCl was used. The simulations were carried out at 298 K. A time step of 4 fs was used with virtual sites for the $M^{pro}$ system, while a 2 fs time step was used for the NBD1 system. A cut-off of 9.5 Å was used for short-range electrostatics and van der Waals interactions in the $M^{pro}$ simulations and a cut-off of 10 Å was used in the NBD1 simulations. The simulation protocol for the $M^{pro}$ system includes 5 ns of simulation with position restraints on heavy atoms, 10 ns of NPT simulation using the Berendsen barostat,[38] and 10 ns NPT simulation using the Parinello-Rahman barostat.[39] The production simulations consisted of eight replicas of 4 $\mu$s using C36m and 5 replicas of 0.8 $\mu$s using ff14SB. The simulation protocol for the NBD1 system is as follows. Position restraints were applied to heavy atoms for 10 ns, followed by 10 ns of NVT, 10 ns NPT using the Berendsen barostat,[38] and 10 ns NPT using the Parinello-Rahman barostat.[39] The production simulations consisted of 5 replicas of 1 $\mu$s each with ff14SB and C36m.

#### 2.15 Elucidating the difference in the rotameric state populations between the ff14SB and C36m ensembles

The distributions of $\chi_1$ and $\chi_2$ rotamers for glutamine residues differ between the C36m and ff14SB simulations (for all three MD systems tested, Fig. 3c and Supplementary Fig. 17). The rotameric state populations in ff14SB simulations are more consistent with the Top8000 dataset[37] than the C36m simulations. The separate analysis of the ff14SB and C36m ensembles (Supplementary Figs. 14 and 15) also pinpoints glutamine side chain dihedrals as the primary cause of inconsistencies between the force fields, which agrees with the combined analysis including the ff94 ensembles.

Here, we describe possible reasons for C36m not capturing the experimental distribution of glutamine rotamers (from Top8000) observed in all systems (Figs. 3c and Supplementary Fig. 17). The side chain dihedral energy parameters in CHARMM36m[40] are the same as in the CHARMM36[41] force field. In the initial optimization of the side chain dihedral parameters in CHARMM36, quantum mechanical (QM) potential energy surfaces were used, followed by additional empirical optimization.[41] In the study in which these QM energy surfaces were obtained,[42] the energy surfaces for amino-acid side chain dihedral angles were converted into probabilities (conforming to a Boltzmann distribution) for a direct comparison with a PDB survey, similar to our comparison to the Top8000 dataset.[37] For the majority of residues, the QM energy surface derived distributions show high consistency with the PDB survey; however, the agreement is low for glutamine, glutamic acid, methionine, as well as lysine, arginine and histidine (Fig. 5 in ref.[42]). Since these QM energy surfaces were used in the optimization of the side chain dihedral parameters in CHARMM36 (and, by extension, CHARMM36m), it is not unexpected that the results of MD simulations using these parameters would show different rotamer populations compared to the Top8000 dataset.

#### 2.16 Markov state models of slow dynamics

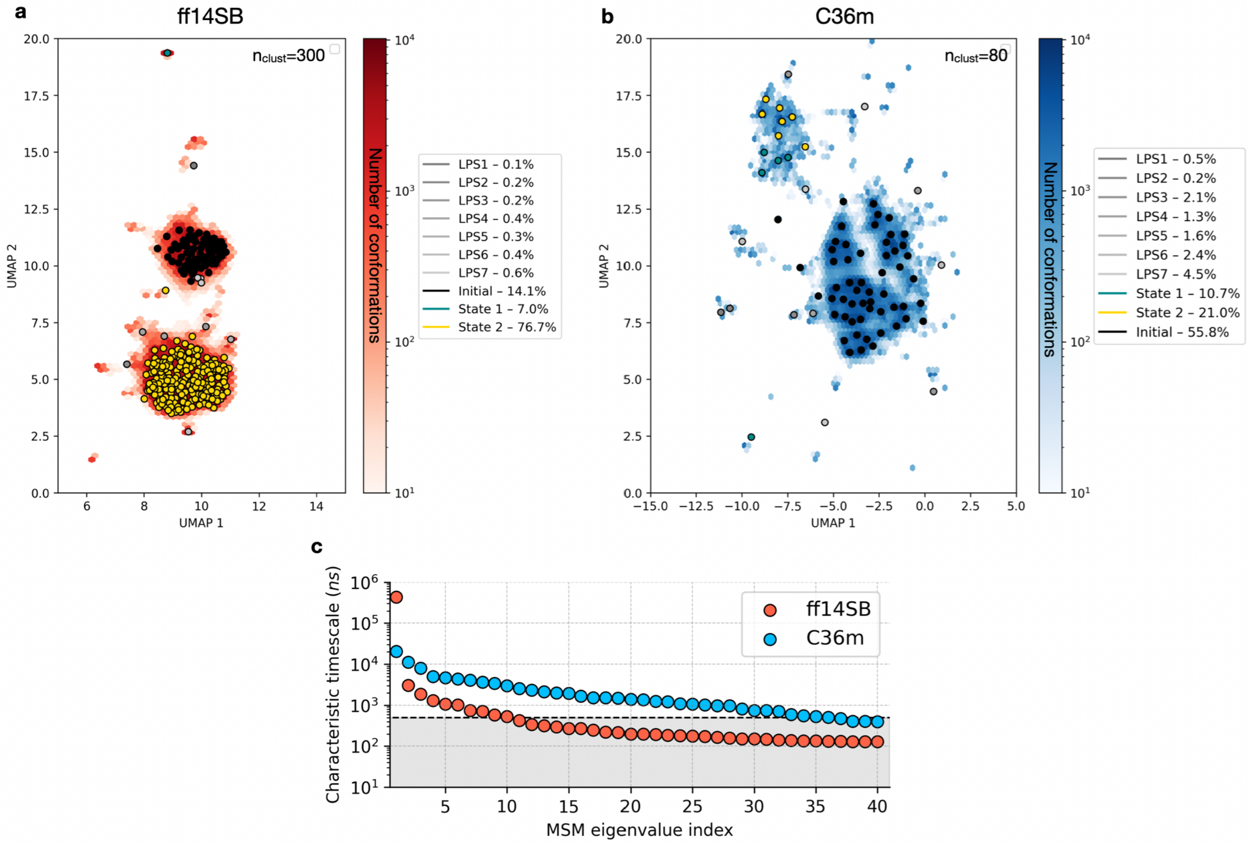

**Supplementary Figure** 18: **Markov state models of slow protein relaxation in the crystal. (a, b)** The two-dimensional UMAP projection of the conformational space, colored by the number of conformations in each hexagon, is shown for the ff14SB and C36m force fields, respectively. The optimum number of microstates (clusters) are: $n_{\text{clust}} = 300$ and $n_{\text{clust}} = 80$ for ff14SB and C36m, respectively. The centroids of these clusters are shown as small circles on each plot. Using kinetic lumping, these microstates were combined into $n = 10$ macrostates to obtain a coarse-grained model describing only the slowest transitions. The legend shows the color and the equilibrium population of each of these macrostates. Macrostates with equilibrium population $< 5\%$ are referred to as low-population states (LPS). The color of each circle indicates the macrostate to which each microstate belongs. **(c)** The top 40 slowest transition timescales (estimated from the MSM eigenvalues) are shown for each MSM in the microstate representation. The timescales of the macrostate models are shown in Fig. 3d.

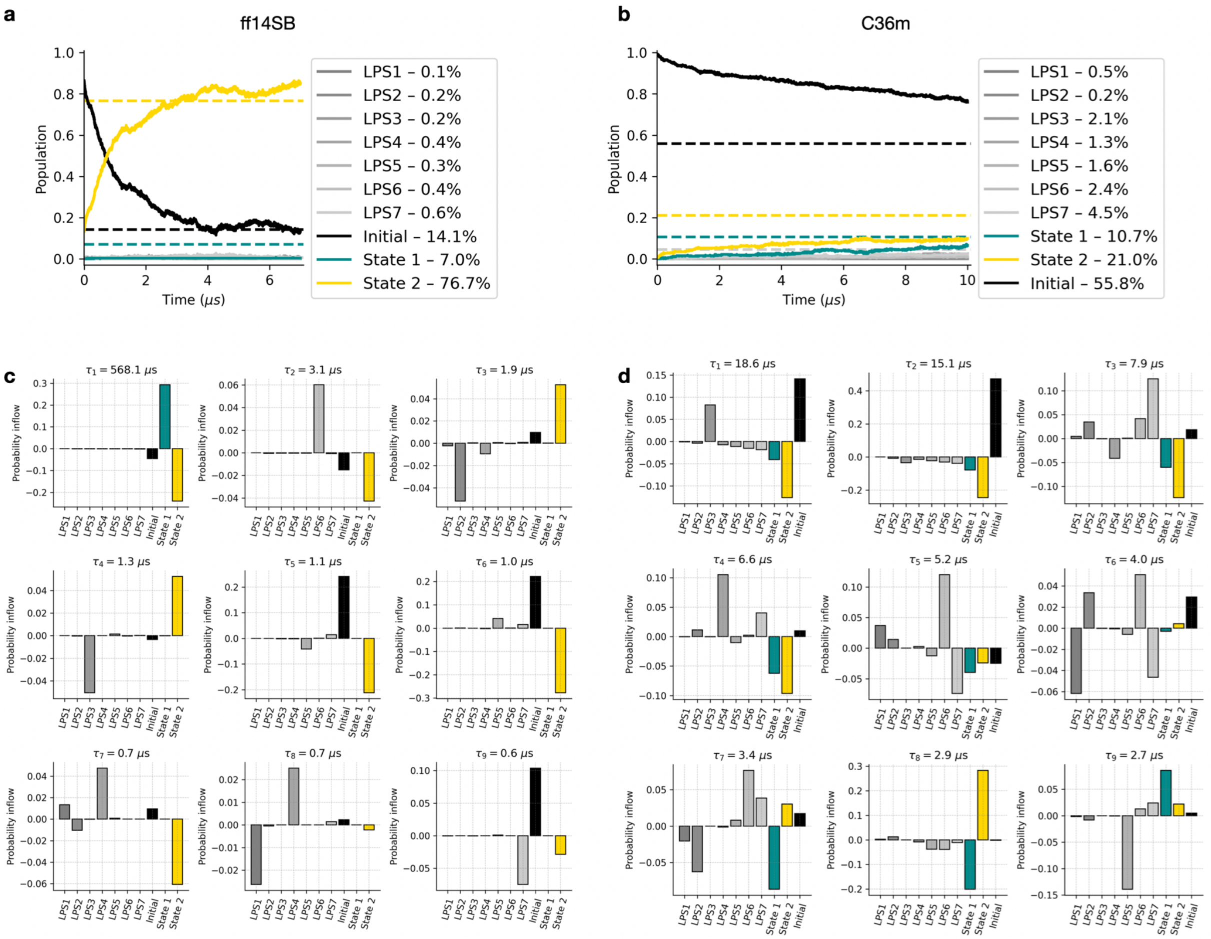

**Supplementary Figure** 19: **MSM state populations and eigenvectors.** **(a, b)** The population of $n = 10$ macrostates as a function of simulation time, for the ff14SB and C36m crystal simulations, respectively. The state population is defined as the number of individual chain trajectories occupying the state at a given time divided by 324 (3 replicas $\times$ 108 chains). Dashed lines show equilibrium populations computed from the MSM transition matrix. States with an equilibrium population of $< 5\%$ are referred to as low-population states (LPS). The structural differences between the three dominant states (initial, state 1 and state 2) in each force field are shown in Supplementary Fig. 20 and visualized in Fig. 3e. **(c, d)** Eigenvectors of the 10-state MSM, for the ff14SB and C36m crystal simulations, respectively. Shown are 9 transitions, with a timescale indicated in the title of each subplot. The conformational transitions happen between the states with high positive and high negative probability inflow.

#### 2.17 Structural differences between Markov states

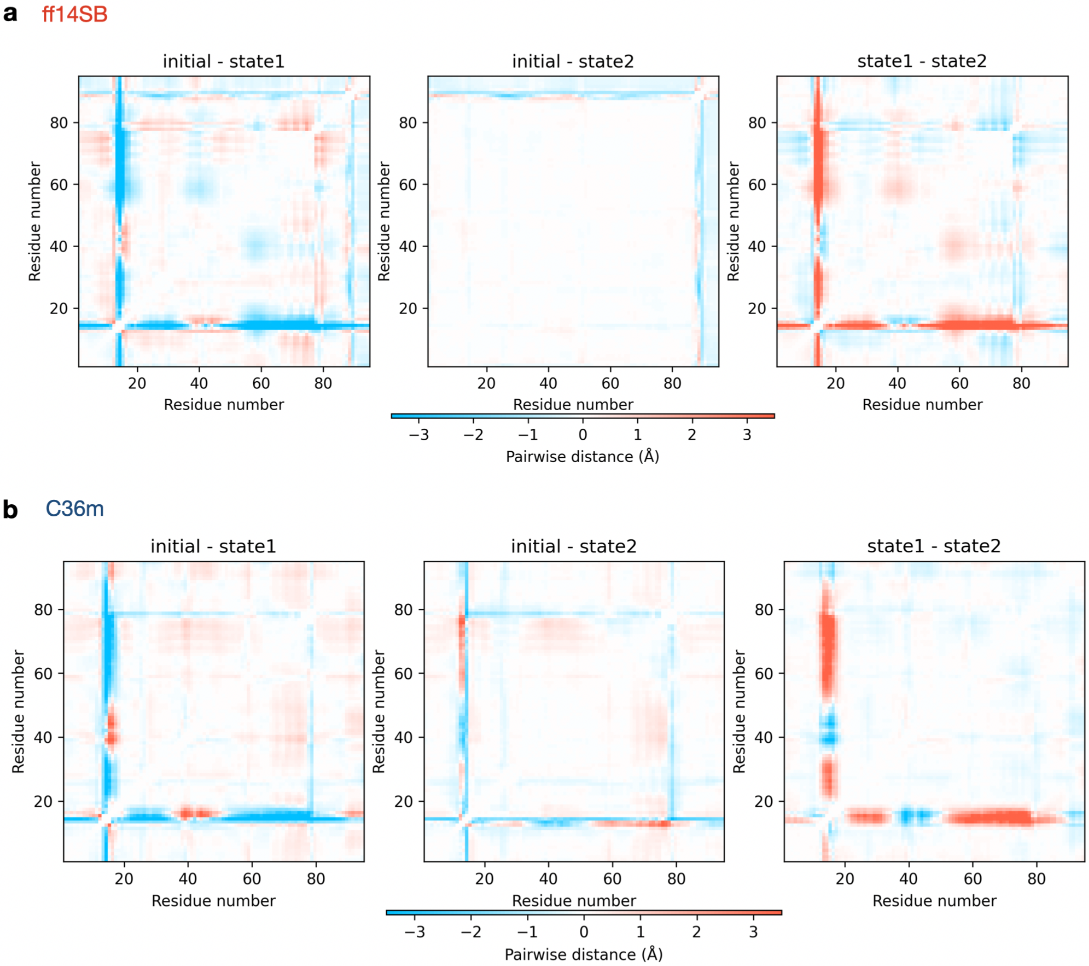

**Supplementary Figure** 20: **Structural differences between dominant states in Markov models.** Matrices of average inter-atomic distances (including only $C\alpha$ atoms) were computed for the initial state, state 1 and state 2 in each ensemble (ff14SB and C36m). The subtraction of pairs of these matrices results in the difference maps shown in **(a)** and **(b)** for ff14SB and C36m, respectively. These difference maps indicate protein regions with the highest deviations between the states, which are illustrated in Fig. 3e of the main text. The population of these three states are shown in Supplementary Fig. 19a-b.

#### 2.18 Distribution of backbone dihedrals

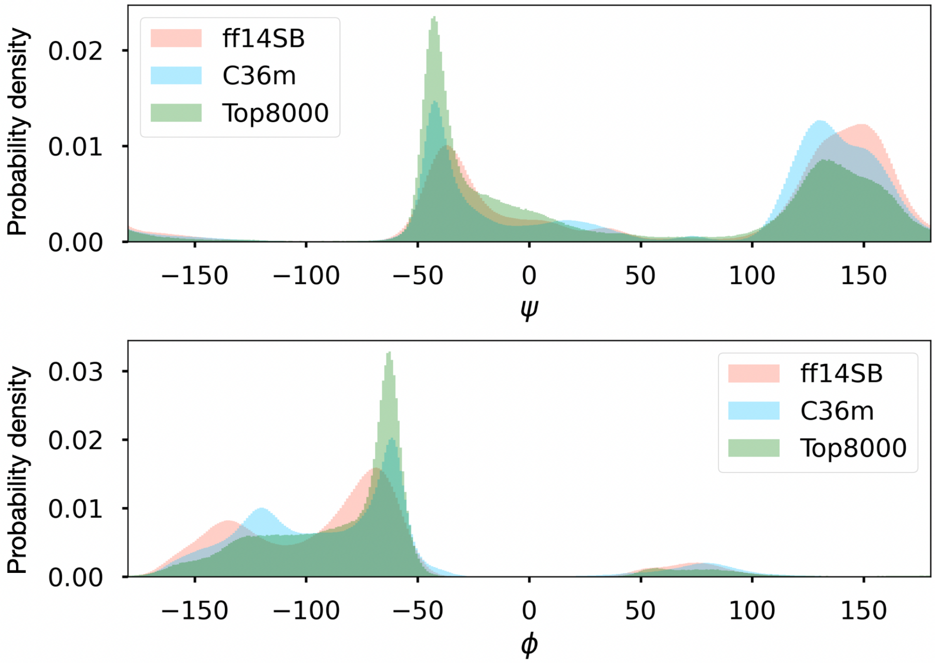

**Supplementary Figure** 21: **The distribution of backbone dihedrals: the contribution of the CMAP correction.** Distribution of Ramachandran angles, $\phi$ and $\psi$, sampled from crystal simulations using ff14SB, C36m, and the Top8000 dataset.[37] Compared to ff14SB, the peaks of the distributions for the C36m dataset are closer to the peaks of the Top8000 distribution. This difference is due in part to the CMAP correction.[43]

#### 2.19 Analysis of the loop conformational states

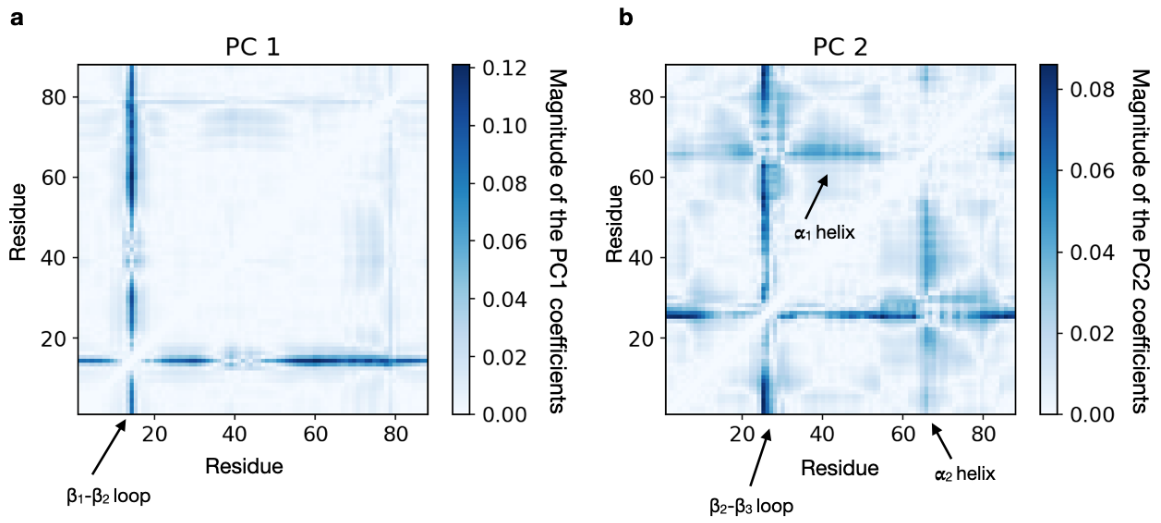

**Supplementary Figure** 22: **Importance of the pairwise** $C\alpha$ **atom distances in principal components 1 and 2**. The PC importance is shown for each $C\alpha$-$C\alpha$ distance for the principal component analysis shown in Fig. 4a of the main text. **(a)** The PC importance is shown for PC 1, which represents 15% of the data variance. The $\beta_1$-$\beta_2$ loop is indicated as the dark blue region around residues 14-16. **(b)** The PC importance is shown for PC 2, which represents 7% of the data variance. The $\beta_2$-$\beta_3$ loop is indicated as a dark blue region around residues 24-26. The dominant conformational change (PC 1) corresponds to the motion of the $\beta_1$-$\beta_2$ loop, while PC 2 represents the much smaller motion of the $\beta_2$-$\beta_3$ loop, and to a lesser extent the $\alpha_1$ helix. The magnitude of the coefficients as well as the percent of data variance in PC 1 are greater than the corresponding values for PC 2, demonstrating a significantly larger contribution of the $\beta_1$-$\beta_2$ loop to the equilibrium motions compared to $\beta_2$-$\beta_3$ loop.

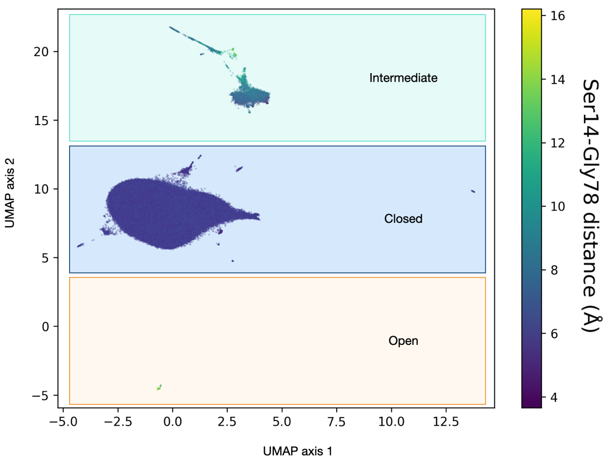

**Supplementary Figure** 23: **The UMAP projection of the equilibrium ff14SB ensemble in the crystal.** The pairwise distances between C$\alpha$ atoms were used as a feature vector (see Methods, Equilibrium protein motions). Structures are colored based on the Ser14-Gly78 distance, defining the three $\beta_1$-$\beta_2$ loop states. Note that the isolated open loop state cluster has a much smaller population than the open and intermediate states. The fluctuation of the $\beta_2$-$\beta_3$ loop, captured by PC 2 (Fig. 4a of the main text), is unrelated to these conformational states and does not correlate with the $\beta_1$-$\beta_2$ loop opening.

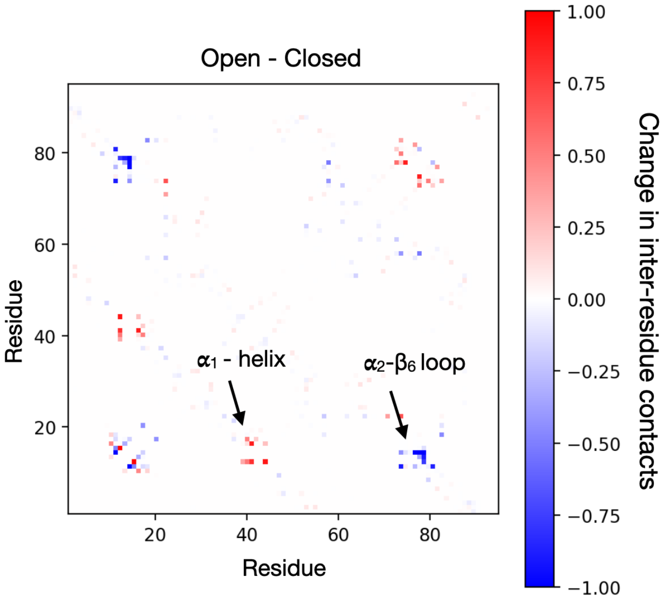

**Supplementary Figure** 24: **Contacts defining the loop states.** The difference contact map between the open and closed loop states shown here was obtained by computing the difference between the open state contact map and the closed state contact map. Contacts that are more probable in the open state are shown in red, while those that are more probable in the closed state are shown in blue.

#### 2.20 Comparison of the protein in solution vs. crystal

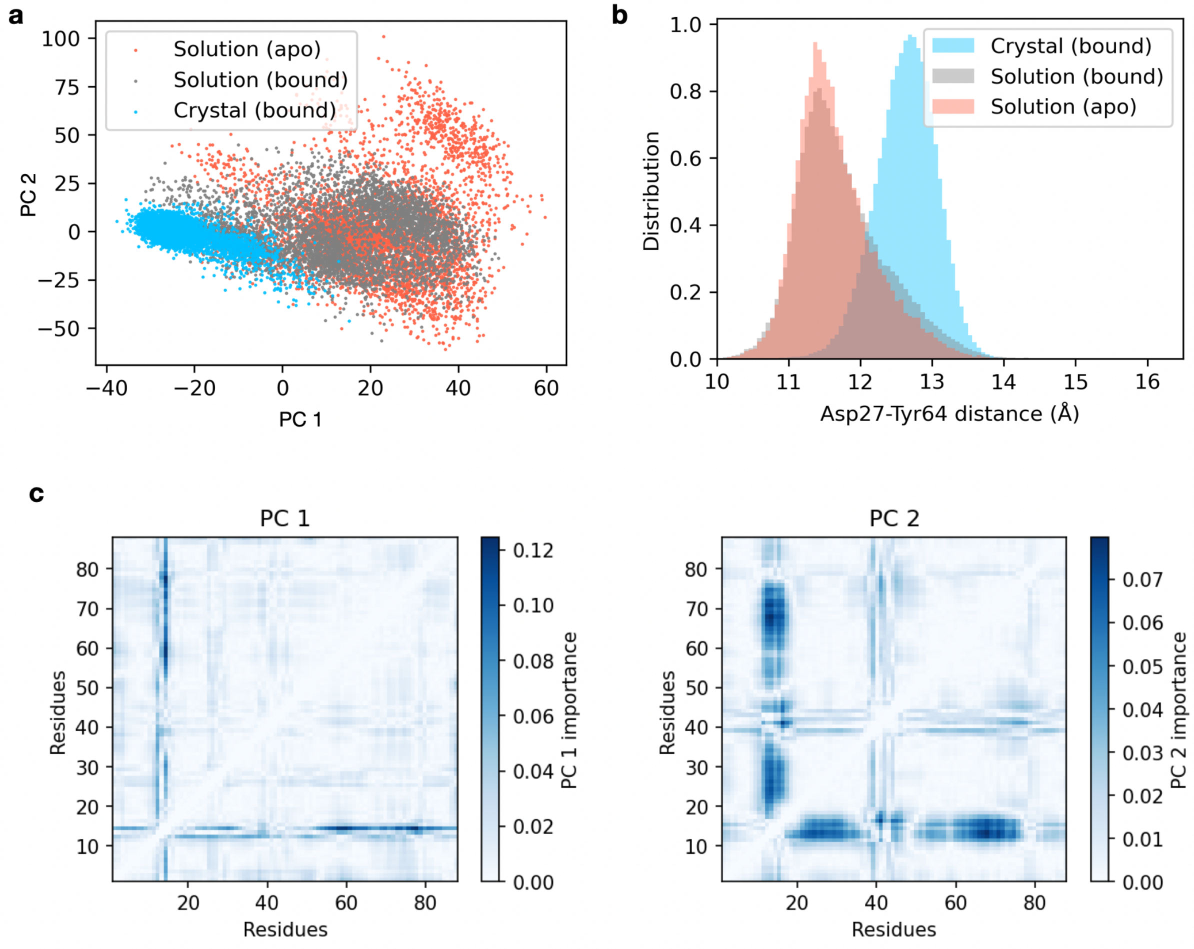

**Supplementary Figure** 25: **Comparison of the protein ensembles in solution vs. crystal.** **(a)** The PCA projection of three ensembles: crystal (ligand-bound), solution (ligand-bound) and solution (ligand-free). Each ensemble was obtained using ff14SB. The same feature vector as in Fig. 4a was used. **(b)** The distribution of the distance between the C$\alpha$ atoms of Asp27 and Tyr64, describing the position of the $\beta_2$-$\beta_3$ loop in the three ensembles. In the crystal structure (PDB ID: 5E11), the Asp27-Tyr64 distance is 12.6 Å. **(c)** The relative PC importance is shown for each $C\alpha$-$C\alpha$ distance for the PCA shown in panel **(a)**. The PC importance is defined as the magnitude of the coefficients of the principal components. PC 1 (24% of variance) corresponds to the motion of the $\beta_1$-$\beta_2$ loop (residues 10-20), while PC 2 (11% of variance) represents the motion in the $\beta_1$-$\beta_2$ loop (residues 10-20) and the $\alpha_1$ helix (residues 39-45).

#### 2.21 Analysis of the strain pseudo-energies

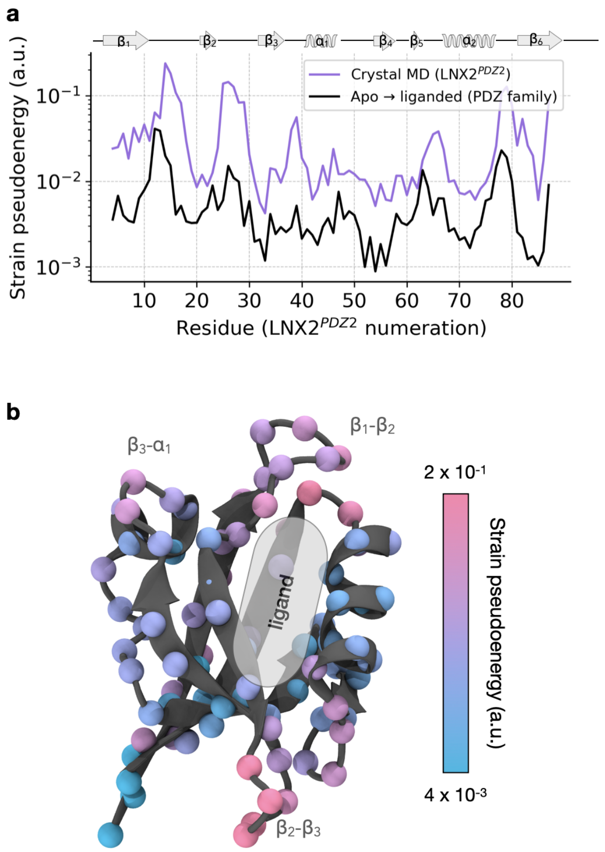

**Supplementary Figure** 26: **Strain profiles**. **(a)** Mean strain pseudo-energies between random pairs of structures from the equilibrium ensemble in the crystal (purple). Median strain pseudo-energies over 10 PDZ domain pairs were mapped on residue numbers of LNX2$^{PDZ2}$ (black). Only C$\alpha$ atoms with matching positions in LNX2$^{PDZ2}$ for each pair of apo- and holo-structures were included in the strain analysis. Strain tensors and strain pseudo-energies were computed using the algorithm described in Mitchell et al.[44] **(b)** Mean strain pseudo-energies between random pairs of structures from the equilibrium ensemble in the crystal (purple line in (a)), mapped onto the PDZ domain structure.

#### 2.22 Analysis of electric-field-induced crystal structures

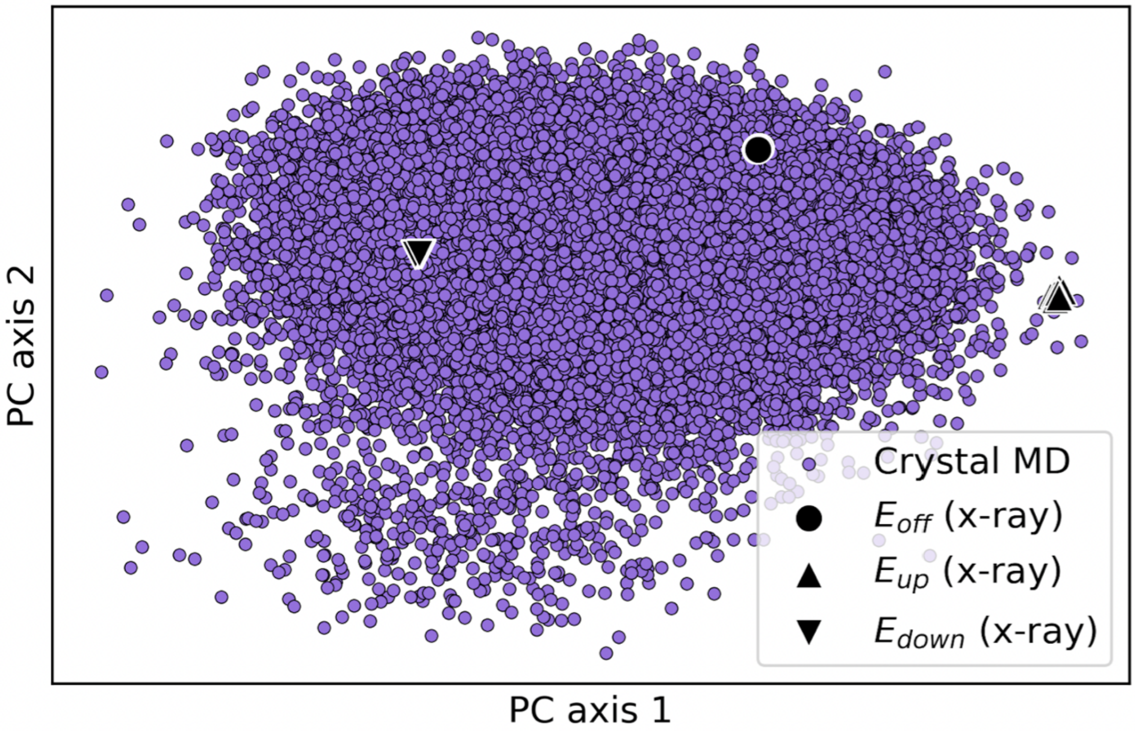

**Supplementary Figure** 27: **Analysis of electric-field-induced crystal structures.** PCA was carried out using crystal structures obtained from the EFX experiment;[45] without an electric field ($E_{\text{off}}$, PDB ID: 5E11) and with an applied electric field ($E_{\text{up}}$ and $E_{\text{down}}$, PDB ID: 5E21). For the experimental structures, all alternative conformations were used and are shown with black markers. In the analysis, the same feature vector as in Fig. 4a of the main text was used. The crystalline ff14SB ensemble projected onto this space is shown in purple.

### 3 Supplementary Methods

#### 3.1 Solvating the crystal lattice

To model the conditions inside the supercell in the simulations, we used the experimental conditions of the crystallization buffer (35 mM of $NaH_2PO_4$ and pH 4.5), reported in the experiment.[45] To mimic the $NaH_2PO_4$ found in the buffer, sodium ($Na^+$) and chloride ($Cl^-$) ions were used. Two approaches were used to model the solvent environment in the crystal. The first approach considers the environment as a simplified solvent, i.e. only containing water and counter-ions. The second approach additionally considers explicit modelling of the crowder molecules found in the crystallization buffer.

**Simplified crystal environment** This approach requires a protocol to carefully add water molecules to the supercell. In the NPT ensemble, the volume of the simulation box changes to preserve the required pressure. In some cases large volume changes may occur if the amount of solvent molecules is insufficient. Hence, when building a protein crystal, the correct number of water molecules must be added to the system to maintain the corresponding experimental volume of the unit cell in the NPT ensemble.[26]

We followed a similar procedure to construct the crystal system as that used by Cerutti and Case.[26] It is important to determine the correct number of water molecules required to preserve the experimental volume in the NPT ensemble.[26] To estimate the optimal number of water molecules required, we performed five independent simulations. Each of these simulations started from a different number of water molecules added to the system. Water molecules were added using GROMACS *gmx solvate*. The range of the number of water molecules to test was determined based on the solvent content of the crystal (e.g. 43% for the PDZ domain). In all cases, crystallographic waters were retained. Each of these systems was simulated as described in Methods, Simulation protocol. The resulting system volume for each system was computed from 100 ns of simulation with anisotropic pressure coupling, then plotted against the number of water molecules added. The optimal number of waters was interpolated from the linear trend (that is, the one producing the correct experimental volume). The system with the optimum number of waters was then used for the pre-production and production runs (refer to Supplementary Fig. 9, plots of volume). The optimal number of waters for each system is provided in Supplementary Table 6.

**Adding explicit crowding agents.** We carried out simulations of the supercell system with explicit crowders. The crystallization buffer (for the crystal used to obtain the structure PDB ID: 5E11) contained polyethylene glycol (PEG) 300 (27-31%) and glycerol (10%).[45] Since the experimental concentration of PEG and glycerol in the crystal is unknown, we aimed to match the concentration in the buffer. To explicitly include crowding agents in the crystal lattice, the following procedure was used.

PEG-282 was chosen for our simulations, as it is most similar in weight to PEG-300 (and PEG-300 does not correspond to any single PEG molecule). The force field parameters were generated using the ACPYPE server[46,47] for Antechamber.[48] To generate a starting pool of crowder conformations, we carried out ten 500 ns simulations of an individual crowder molecule (PEG or glycerol) in a box of water at 300 K. From the resulting pool, randomly selected conformations of 362 PEG molecules and 428 glycerol molecules were inserted into the supercell with crystallographic water molecules and water molecules added to match the experimental volume (Supplementary Table 6). On average, there are 12 and 16 molecules of PEG and glycerol per unit cell, respectively.

#### 3.2 Simulations of the PDZ domain in solution

Two types of simulations of the PDZ domain in solution were carried out: in the ligand-free and ligand-bound form.

**Ligand-free form**. Solution simulations in a ligand-free form were performed using ff14SB and C36m force fields. The simulation protocol matched the one used for the crystal simulations. Each simulation replica was initialized from the same atomic coordinates and randomly seeded velocities. Ten simulation replicas were carried out for 3 $\mu$s for ff14SB and 4.5$\mu$s for C36m.

We analyzed the heavy-atom RMSD as a function of time (Supplementary Fig. 28a). The C-terminal tail was excluded from the analysis due to its high mobility in solution. In the crystal, the C-terminal motif (residues 92-95) is found in the binding site of an adjacent chain. However, in solution, this segment is unbound and, therefore, has increased conformational freedom. Equilibration times are estimated to be 1 $\mu$s for ff14SB and 3 $\mu$s for C36m (Supplementary Fig. 28a). The mean RMSD value does not exceed 2.5 Å in the ff14SB simulations, whereas it reaches 4.0 Å in the C36m simulations. The major difference between the C36m ensemble and the crystal structure (PDB ID: 5E11) is in the $\alpha 1$ helix (Supplementary Fig. 28b). The ff14SB force field also yields a more structurally consistent ensemble than C36m, as seen in the lower variance of RMSD across $n = 10$ replicas (shaded envelope in Supplementary Fig. 28a). Taken together, these results suggest that ff14SB is more consistent in preserving the crystal structure when compared to C36m, not only in the crystal (Figs. 1b, 1c, 2a, and 2b), but also in solution.

**Ligand-bound form.** To investigate the functional relevance of protein motions in the crystal, we needed to test whether these motions are affected by ligand binding. We therefore simulated the same PDZ domain in a ligand-bound form, where the 4-residue ligand-peptide (residues 92-95 of the C-terminal tail) was bound to the active site of the protein. Residue 92 (the N-terminus of the peptide) was methylated to neutralize the charge and to avoid electrostatic interaction (i.e. to maintain a similar interaction as with the full-

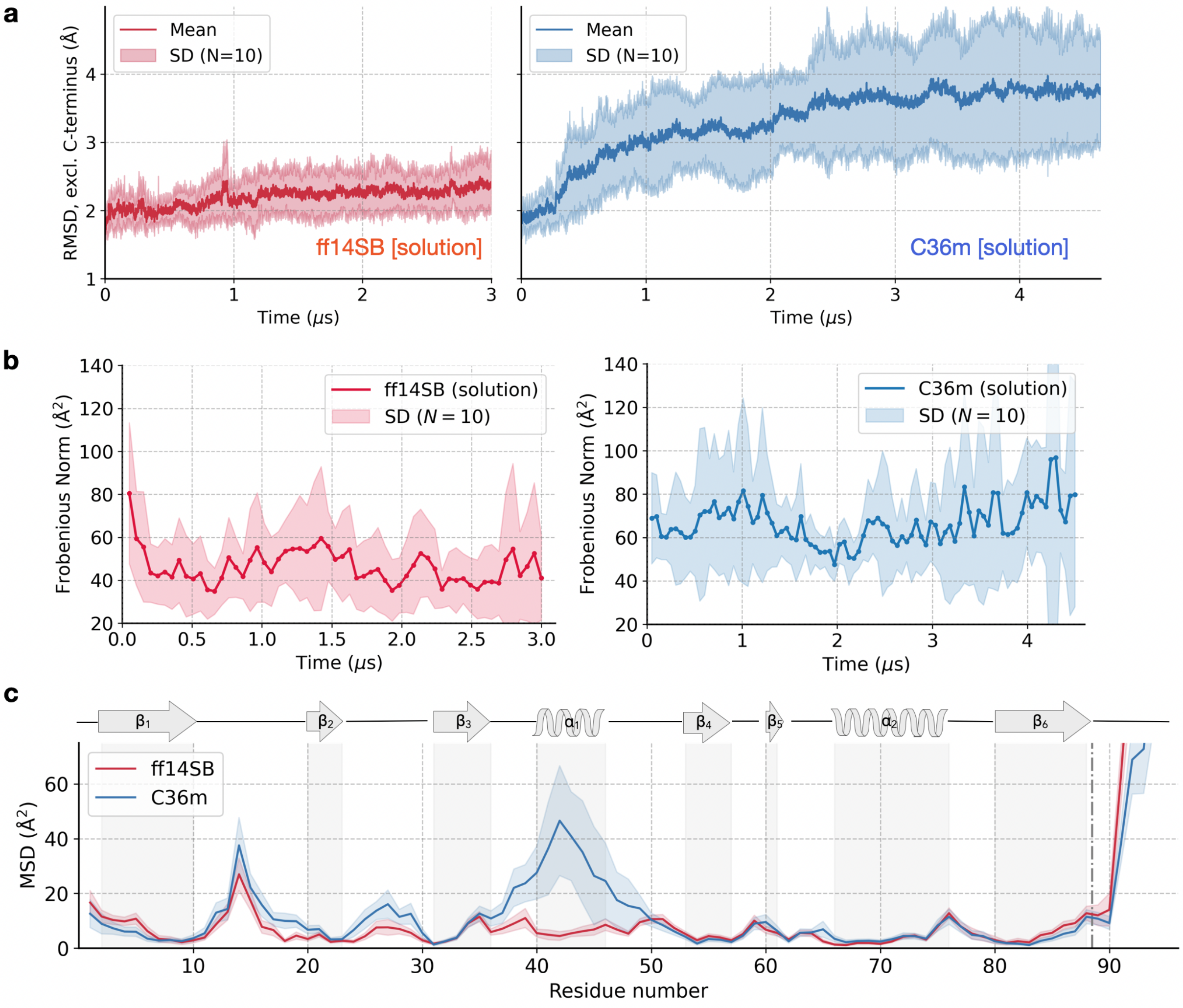

**Supplementary Figure** 28: **Analysis of solution simulations in the ligand-free form**. **(a)** The RMSD (including all heavy atoms in residues 1-89) from the crystal structure (PDB ID: 5E11) as a function of time using the ff14SB (left) and C36m (right) force fields. The shaded envelope represents the mean ± standard deviation over $n = 10$ replicas. **(b)** Convergence of the atomic covariance matrix, calculated as the Frobenius norm (similar to Supplementary Fig. 3) as a function of time for the solution simulations using the ff14SB (left) and C36m (right) force fields. The shaded envelope represents the mean ± standard error over $n = 10$ replicas. **(c)** The average MSD of C$\alpha$ atoms relative to the crystal structure, with the shaded envelope representing the mean ± standard deviation over $n = 10$ replicas.

length PDZ domain). For this system, we carried out nine simulations of 2 $\mu$s each using the Amber ff14SB force field and the TIP3P water model. A comparison of the RMSD to the crystal structure (PDB ID: 5E11) between ligand-free and ligand-bound systems are shown in Supplementary Fig. 29, where both systems reach a plateau.

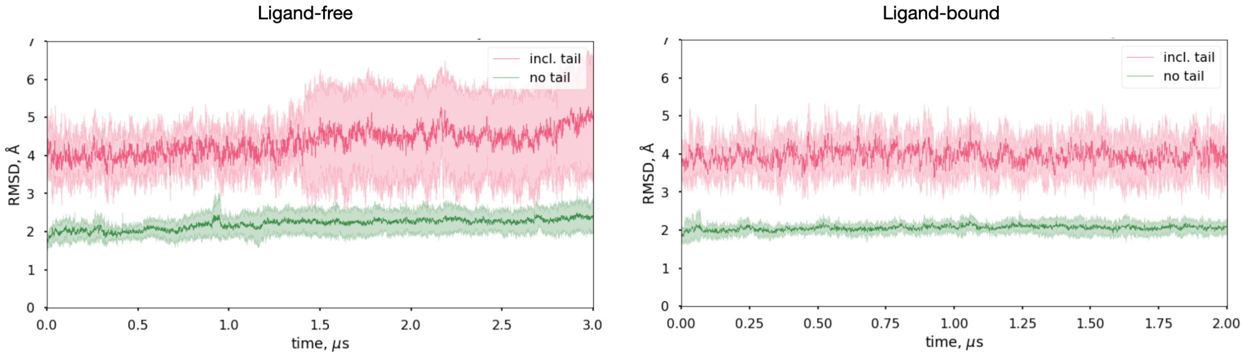

**Supplementary Figure** 29: **Comparing RMSD between apo and bound solution simulations using ff14SB**. The heavy-atom RMSD compared to the crystal structure (PDB ID: 5E11) as a function of time. The shaded envelope represents the mean ± standard deviation over $n = 10$ replicas (ligand-free, left) and over $n = 9$ replicas (ligand-bound, right). The RMSD of the entire protein is shown in red, and the RMSD without the flexible C-terminal tail (residues 89-95) is shown in green.

#### 3.3 Simulation Systems

**Supplementary Table** 6: Number of atoms in each simulation system.

| Simulation system | $H_2O$ | $Na^+$ | $Cl^-$ | # of atoms |
|---|---|---|---|---|
| Amber ff14SB (supercell) | 33,555 | 159 | 51 | 259,959 |
| CHARMM36m (supercell) | 34,186 | 159 | 51 | 261,852 |
| Amber ff94 (supercell) | 33,771 | 159 | 51 | 260,607 |
| Amber ff14SB (supercell + crowd.) | 26,952 | 159 | 51 | 262,432 |
| Amber ff14SB (solution, apo) | 11,863 | 9 | 8 | 37,079 |
| CHARMM36m (solution, apo) | 11,863 | 9 | 8 | 37,079 |
| Amber ff14SB (solution, bound) | 11,864 | 12 | 8 | 37,160 |

#### 3.4 Analysis

Multiple metrics to quantify structural similarity and convergence of simulations were used: RMSD, fraction of native contacts (Q), per-residue mean squared deviation (MSD), and the covariance matrix.

**RMSD and MSD.** In the calculation of RMSD, we only considered deviations of heavy atoms between the simulated and experimental structures. Instead of comparing the simulated proteins to a single reference structure, the RMSD of a given simulation structure was referenced against 16 possible alternative conformations ($AAAA, AAAB, AABA, ..., BBBB$, see below for details) found in the PDB file, only considering the minimum RMSD:

$$\mathrm{RMSD}_{\mathrm{min}}(t) = \min\left(\mathrm{RMSD}_1(t), \mathrm{RMSD}_2(t), ..., \mathrm{RMSD}_{16}(t)\right). \quad (1)$$

The mean-squared deviations, $\delta_i^2$, for each C$\alpha$ atom $i$ elucidate the regions of the protein which deviate the most from the crystal structure (PDB ID: 5E11). MSDs $\delta_i^2$ were computed by aligning each protein chain in the supercell to the reference structure (i.e. the alternate conformation having the minimum RMSD to the chain of interest) averaged over time and chains:

$$\delta_i^2 = \frac{1}{N_{\mathrm{chains}}} \sum_{j=1}^{N_{\mathrm{chains}}} \langle (\mathbf{x}_{ij} - \mathbf{x}_i^{ref})^2 \rangle, \quad (2)$$

where $\langle . \rangle$ denotes a time-average, $\mathbf{x}_{ij}$ is the vector position of an atom $i$ in the protein chain $j$, $\mathbf{x}_i^{ref}$ is the position of the atom in the reference structure, and $N_{\mathrm{chains}} = 108$ for the $3 \times 3 \times 3$ supercell.

**Alternative conformations.** The experimental structure (PDB ID: 5E11) of the PDZ domain includes four regions of the protein that have two alternative conformations (A and B) in the crystal. Region 1 corresponds to residue 17, region 2 includes residues 34 to 36, region 3 includes residues 48 to 51, and region 4 includes residues 80 to 85. Since each of these four regions has two conformations, 16 independent structures (which we refer to as states) ($N_{\text{Conf.}}^{M_{\text{Reg.}}} = 2^4$) were constructed. These 16 structures are referred to as $AAAA, AAAB, AABA, ..., BBBB$, where each letter signifies the alternative conformation at the position of the corresponding protein region. The RMSD computed between all pairs of 16 states (Supplementary Fig. 30) indicates that the largest differences between alternative conformations occur due to the fourth region (e.g. comparing AAAB and AAAA). The maximum RMSD between two alternative states in the crystal is 0.52 Å.

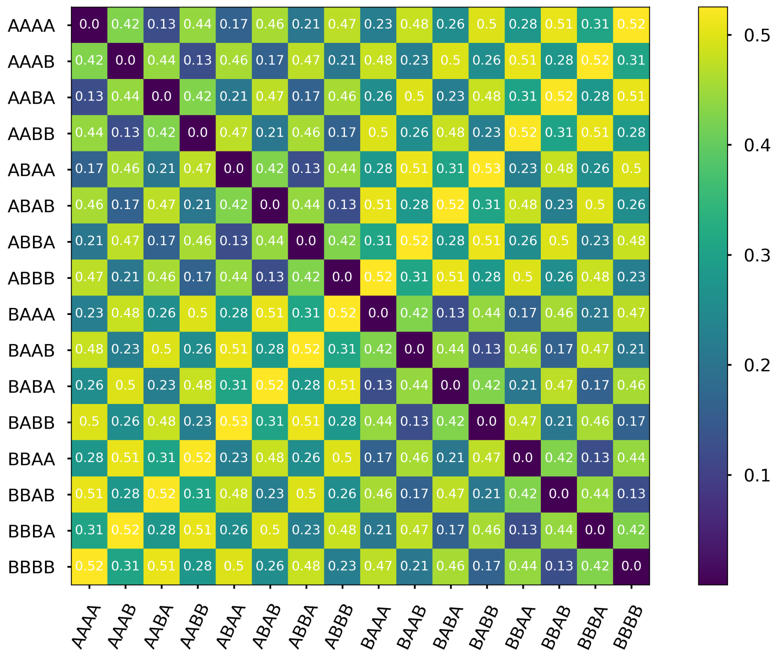

**Supplementary Figure** 30: **Deviation between alternative conformations.** Shown are the RMSD values (in Å) between all pairs of 16 alternative conformational states ($AAAA, AAAB, AABA, ..., BBBB$) in PDB ID: 5E11. RMSD is computed using only heavy atoms in this analysis.

**Covariance matrix.** The Python module *MDAnalysis.PCA* was used to compute covariance matrices (Supplementary Figs. 3, 6 and 28b). Each trajectory was aligned to the initial frame, minimizing the RMSD for $C\alpha$ atoms. The resulting covariance matrices, denoted as $\text{cov}_{\beta\gamma}$ and representing pairs of coordinates $\beta, \gamma$ for $C\alpha$ atoms, were computed for individual chains within the supercell using 100-ns windows with 1000 frames per window. The dimensions of each covariance matrix were $285 \times 285$, where 285 is the product of the number of $C\alpha$ atoms (95) and the number of Cartesian coordinates (3) in each row/column.

To assess convergence, the Frobenius norm distance between covariance matrices $\mathrm{cov}_{\beta\gamma}$ at time $t$ and $t + 100$ ns was calculated. This distance, averaged over $n = 108$ chains, was plotted against time $t$ as a way to assess the convergence of the simulations (Supplementary Fig. 3).

To assess the type of atomic motions in the crystal (Supplementary Fig. 6), the covariance matrix $\mathrm{cov}_{ij}$ and distance matrix $d_{ij}$ between pairs of $C\alpha$ atoms $i, j$ were computed for the entire supercell, following the protocol described by Wych et al.[22] Initially, the covariance matrix for individual atomic coordinates, $\mathrm{cov}_{\beta\gamma}$, was calculated with size $30,780 \times 30,780$, where $30,780 = 108$ chains $\times 95$ atoms $\times 3$ coordinates. Next, $\mathrm{cov}_{ij}$ was created by computing a trace of each $3\times3$ submatrix of $\mathrm{cov}_{\beta\gamma}$ to represent covariance between individual atoms. The computation utilized the last 100 ns (1000 frames) of each simulation replica, with subsequent averaging over three replicas. The distance matrix $r_{ij}$ underwent two-tier averaging, first over 1000 frames and then over the three replicas. Both matrices ($\mathrm{cov}_{ij}$ and $d_{ij}$) share dimensions of $10,260 \times 10,260$, where $10,260 = 108$ chains $\times 95$ atoms. Subsequently, the values of the distance matrix were distributed into 60 bins ranging from 0 to 120 Å with a bin size of 2 Å. To plot atomic covariance against distance in a crystal, the covariance values $\mathrm{cov}_{ij}$ for pairs of atoms within each $r_{ij}$ bin were averaged. The resulting curve, denoted as $\mathrm{cov}_{ij}$ vs. $r_{ij}$, was fitted to an exponential function $ae^{-r_{ij}/\gamma} + b$. The error estimates for each fitted parameter were calculated from the variance values provided by the *scipy.optimize.curve_fit* function in Python and represent 95% confidence intervals.

**Contacts.** Residue contacts were defined as follows: a contact between two residues is counted if the distance between any pair of heavy atoms is less than 5.5 Å. For the contact map difference (Fig. 10), we excluded all transient contacts with propensity $< 0.5$ from the analysis. The fraction of native intra-protein contacts, $Q$, is the number of common contacts that are present both in the experimental structure and in a structure from the simulation, divided by the total number of contacts in the crystal structure (PDB ID: 5E11).

**B-factors.** Experimental isotropic B-factors were obtained from the crystal structure

(PDB ID: 5E11).[45] Computed B-factors are related to the root mean square fluctuation (RMSF) of atomic positions:

$$B_i = \frac{8\pi^2}{3}\rho_i^2. \tag{3}$$

RMSF of an atom $\rho_i = \sqrt{\langle(\mathbf{x}_i - \langle\mathbf{x}_i\rangle)^2\rangle}$ where $\langle.\rangle$ denote a time average, and $\mathbf{x}_i$ is the vector-position of atom $i$. In the case of a protein supercell simulation, we used two methods to estimate RMSF, depending on the alignment procedure, following an approach similar to Cerutti et al.[13] In the first method, referred to as $B_{\text{lattice}}$, we removed the center of mass drift of the whole supercell and computed RMSF for every atom in the supercell. We then averaged atomic RMSFs $\rho_i^2$ over all 108 copies. In the second method, which we denote $B_{\text{chain}}$, the 108 chains were isolated and aligned to a reference structure. The atomic RMSFs, $\rho_i^2$, were calculated and averaged over all 108 chains. This method removes rotations and translations of individual chains within a lattice. The equilibrated portion (the last 100 ns) of each trajectory were used for computing RMSFs. Final B-factor values (Fig. 2c) were obtained by averaging over 3 simulation replicas.

**Average MD structures.** The average MD structure for each force field was computed from the supercell simulations (3 replicas $\times$ 108 chains per replica for a total of 324 individual trajectories). The structure at each frame was aligned to the crystal structure using backbone atoms. Using these trajectories, 324 time-averaged structures were computed, which were then averaged to produce a single average structure. It should be noted that the resulting average structure is non-physical, because realistic atomic geometries were not enforced. To produce a "physical" average structure (shown in Fig. 3b), the entire simulation ensemble was then scanned to identify the conformation that is the most similar to the average, i.e. the structure with the minimum heavy-atom RMSD (0.70 Å for C36m and 0.72 Å for ff14SB).

**Rotameric states of side chain dihedrals.** The rotameric states are defined as follows: gauche+ ($g+$) is between 0° and 120°, trans ($t$) is between 120° and 240°, gauche− ($g-$) is between 240° and 360°.

**Markov state models**. In constructing MSMs for the crystal simulations, we chose a lag time for both ff14SB and C36m data sets based on the convergence of the slowest characteristic timescales for microstate transitions as a function of lag time (Supplementary Fig. 31). We note that the optimum lag time did not depend on the number of clusters used as soon as there were more than 50 clusters. Once the lag time was fixed ($\tau = 500$ ns), we utilized a 10-fold cross-validation (by splitting a dataset into the train and test sets) to find the optimum number of microstates (clusters) based on generalized matrix Rayleigh quotient score (GMRQ, Supplementary Fig. 31c,d). We picked the value of parameter $n_{\mathrm{clust}}$ which gave the maximum median score on the test data set across all 10 folds. Using a long lag time (500 ns) and a large number of microstates (from 80 to 300) reduces the bias in the estimated MSMs.[49]

**Structural analysis of related PDZ domains.** Ten pairs of high-resolution (< 2 Å) crystal structures of PDZ domains with and without ligand were used: E3-LNX$^{PDZ}$: 3VQG vs. 3VQF; PSD-95$^{PDZ3}$: 1BFE and 6QJJ vs. 1BE9; PTP-1E$^{PDZ2}$: 3LNX vs. 3LNY; GRIP-1$^{PDZ6}$: 1N7E vs. 1N7F; SAP97$^{PDZ2}$: 2AWU vs. 2AWW; PALS-1$^{PDZ}$: 4UU6 vs. 4UU5; Erbin$^{PDZ}$: 2H3L vs. 1MFG; Dishevelled$^{PDZ}$: 2F0A vs. 1L6O; PDZK-1$^{PDZ3}$: 3R68 vs. 3R69. These PDZ domains were chosen because they have the highest sequence similarity to LNX2$^{PDZ2}$ and they have both apo and ligand-bound crystal structures available. Apo and liganded structures were first aligned to each other in PyMOL[50] using *align* for C$\alpha$ atoms, then aligned to the crystal structure of LNX2$^{PDZ2}$ (PDB ID 5E11) using *super*. The median (over PDZ domains) ligand-induced deviation of $C\alpha$ atoms (Fig. 5a, black line) were computed as described in Fig. 5e of Hekstra et al.[45] Only residues with matching positions in LNX2$^{PDZ2}$ for each pair of apo and bound structures were included in the analysis.

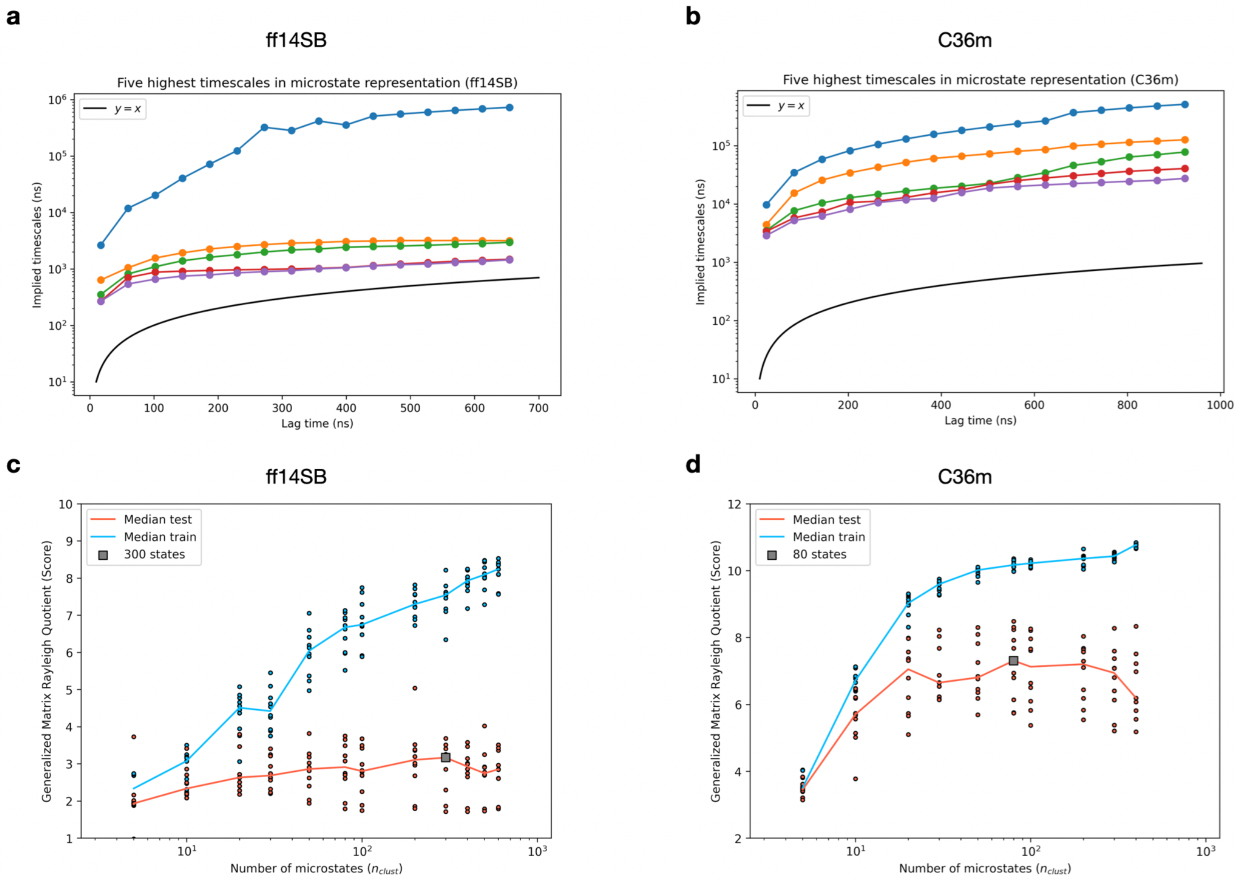

**Supplementary Figure** 31: **Markov state model optimization: lag time ($\tau$) and number of microstates ($n_{\mathbf{clust}}$).** The slowest transition timescales between microstates as a function of lag time for the **(a)** ff14SB and **(b)** C36m crystal simulations. After a lag time was chosen based on convergence of the plot ($\tau = 500$ ns), each model was optimized using the generalized matrix Rayleigh quotient (GMRQ) score to find the optimum number of microstates. A 10-fold cross-validation was employed to determine the value of the parameter $n_{\mathrm{clust}}$ that results in the highest GMRQ score on the test set. The median score over the ten samples was used. These results are shown for the **(c)** ff14SB and **(d)** C36m force fields, with the optimum number of microstates indicated in gray. Details of the microstate definitions are provided in the Methods section of the main text.

### Supplementary References

(1) Gunsteren, W. F. V.; Karplus, M. Protein dynamics in solution and in a crystalline environment: a molecular dynamics study. *Biochemistry* **1982**, *21*, 2259–2274, DOI: 10.1021/bi00539a001.

(2) van Gunsteren, W. F.; Berendsen, H. J.; Hermans, J.; Hol, W. G.; Postma, J. P. Computer simulation of the dynamics of hydrated protein crystals and its comparison with x-ray data. *Proceedings of the National Academy of Sciences* **1983**, *80*, 4315–4319, DOI: 10.1073/pnas.80.14.4315.

(3) Berendsen, H.; van Gunsteren, W.; Zwinderman, H.; Geurtsen, R. Simulations of Proteins in Water. *Annals of the New York Academy of Sciences* **1986**, *482*, 269–286, DOI: 10.1111/j.1749-6632.1986.tb20961.x.

(4) Avbelj, F.; Moult, J.; Kitson, D. H.; James, M. N. G.; Hagler, A. T. Molecular dynamics study of the structure and dynamics of a protein molecule in a crystalline ionic environment, Streptomyces griseus protease A. *Biochemistry* **1990**, *29*, 8658–8676, DOI: 10.1021/bi00489a023.

(5) García, A. E.; Blumenfeld, R.; Hummer, G.; Krumhansl, J. A. Multi-basin dynamics of a protein in a crystal environment. *Physica D: Nonlinear Phenomena* **1997**, *107*, 225–239, DOI: 10.1016/s0167-2789(97)00090-0.

(6) Ceccarelli, M.; Marchi, M. Simulation of a Protein Crystal at Constant Pressure. *The Journal of Physical Chemistry B* **1997**, *101*, 2105–2108, DOI: 10.1021/jp9701810.

(7) Héry, S.; Genest, D.; Smith, J. C. Fluctuation and Correlation in Crystalline Lysozyme. *Journal of Chemical Information and Computer Sciences* **1997**, *37*, 1011–1017, DOI: 10.1021/ci970234a.

(8) Stocker, U.; Spiegel, K.; van Gunsteren, W. On the similarity of properties in solution or in the crystalline state: A molecular dynamics study of hen lysozyme. *Journal of Biomolecular NMR* **2000**, *18*, 1–12, DOI: 10.1023/a:1008379605403.

(9) Meinhold, L.; Smith, J. C. Fluctuations and Correlations in Crystalline Protein Dynamics: A Simulation Analysis of Staphylococcal Nuclease. *Biophysical Journal* **2005**, *88*, 2554–2563, DOI: 10.1529/biophysj.104.056101.

(10) Joti, Y.; Nakagawa, H.; Kataoka, M.; Kitao, A. Hydration-Dependent Protein Dynamics Revealed by Molecular Dynamics Simulation of Crystalline Staphylococcal Nuclease. *The Journal of Physical Chemistry B* **2008**, *112*, 3522–3528, DOI: 10.1021/jp710039p.

(11) Cerutti, D. S.; Trong, I. L.; Stenkamp, R. E.; Lybrand, T. P. Simulations of a Protein Crystal: Explicit Treatment of Crystallization Conditions Links Theory and Experiment in the Streptavidin-Biotin Complex. *Biochemistry* **2008**, *47*, 12065–12077, DOI: 10.1021/bi800894u.

(12) Hu, Z.; Jiang, J. Assessment of biomolecular force fields for molecular dynamics simulations in a protein crystal. *Journal of Computational Chemistry* **2010**, *31*, 371–380, DOI: 10.1002/jcc.21330.

(13) Cerutti, D. S.; Freddolino, P. L.; Duke, R. E.; Case, D. A. Simulations of a Protein Crystal with a High Resolution X-ray Structure: Evaluation of Force Fields and Water Models. *The Journal of Physical Chemistry B* **2010**, *114*, 12811–12824, DOI: 10.1021/jp105813j.

(14) Janowski, P. A.; Cerutti, D. S.; Holton, J.; Case, D. A. Peptide Crystal Simulations Reveal Hidden Dynamics. *Journal of the American Chemical Society* **2013**, *135*, 7938–7948, DOI: 10.1021/ja401382y.

(15) Wall, M. E.; Benschoten, A. H. V.; Sauter, N. K.; Adams, P. D.; Fraser, J. S.; Terwilliger, T. C. Conformational dynamics of a crystalline protein from microsecond-scale

molecular dynamics simulations and diffuse X-ray scattering. *Proceedings of the National Academy of Sciences* **2014**, *111*, 17887–17892, DOI: 10.1073/pnas.1416744111.

(16) Xue, Y.; Skrynnikov, N. R. Ensemble MD simulations restrained via crystallographic data: Accurate structure leads to accurate dynamics. *Protein Science* **2014**, *23*, 488–507, DOI: 10.1002/pro.2433.

(17) Li, Y.; Zhang, J. Z. H.; Mei, Y. Molecular Dynamics Simulation of Protein Crystal with Polarized Protein-Specific Force Field. *The Journal of Physical Chemistry B* **2014**, *118*, 12326–12335, DOI: 10.1021/jp503972j.

(18) Kuzmanic, A.; Pannu, N. S.; Zagrovic, B. X-ray refinement significantly underestimates the level of microscopic heterogeneity in biomolecular crystals. *Nature Communications* **2014**, *5*, 3220, DOI: 10.1038/ncomms4220.

(19) Janowski, P. A.; Liu, C.; Deckman, J.; Case, D. A. Molecular dynamics simulation of triclinic lysozyme in a crystal lattice. *Protein Science* **2015**, *25*, 87–102, DOI: 10.1002/pro.2713.

(20) Wall, M. E. Internal protein motions in molecular-dynamics simulations of Bragg and diffuse X-ray scattering. *IUCrJ* **2018**, *5*, 172–181, DOI: 10.1107/s2052252518000519.

(21) Zhu, T.; Wu, C.; Song, J.; Reimers, J. R.; Li, Y. Polarization effect within a protein crystal: A molecular dynamics simulation study. *Chemical Physics Letters* **2018**, *706*, 303–307, DOI: 10.1016/j.cplett.2018.06.018.

(22) Wych, D. C.; Fraser, J. S.; Mobley, D. L.; Wall, M. E. Liquid-like and rigid-body motions in molecular-dynamics simulations of a crystalline protein. *Structural Dynamics* **2019**, *6*, 064704, DOI: 10.1063/1.5132692.

(23) Meisburger, S. P.; Case, D. A.; Ando, N. Diffuse X-ray scattering from corre-

lated motions in a protein crystal. *Nature Communications* **2020**, *11*, 1271, DOI: 10.1038/s41467-020-14933-6.

(24) Wych, D. C.; Aoto, P. C.; Vu, L.; Wolff, A. M.; Mobley, D. L.; Fraser, J. S.; Taylor, S. S.; Wall, M. E. Molecular-dynamics simulation methods for macromolecular crystallography. *Acta Crystallographica Section D Structural Biology* **2023**, *79*, 50–65, DOI: 10.1107/s2059798322011871.

(25) Jo, S.; Kim, T.; Iyer, V. G.; Im, W. CHARMM-GUI: A web-based graphical user interface for CHARMM. *Journal of Computational Chemistry* **2008**, *29*, 1859–1865, DOI: 10.1002/jcc.20945.

(26) Cerutti, D. S.; Case, D. A. Molecular dynamics simulations of macromolecular crystals. *WIREs Computational Molecular Science* **2018**, *9*, e1402, DOI: 10.1002/wcms.1402.

(27) Doyle, D. A.; Lee, A.; Lewis, J.; Kim, E.; Sheng, M.; MacKinnon, R. Crystal structures of a complexed and peptide-free membrane protein-binding domain: molecular basis of peptide recognition by PDZ. *Cell* **1996**, *85*, 1067–1076, DOI: 10.1016/s0092-8674(00)81307-0.

(28) Fuentes, E. J.; Der, C. J.; Lee, A. L. Ligand-dependent Dynamics and Intramolecular Signaling in a PDZ Domain. *Journal of Molecular Biology* **2004**, *335*, 1105–1115, DOI: 10.1016/j.jmb.2003.11.010.

(29) Lockless, S. W.; Ranganathan, R. Evolutionarily Conserved Pathways of Energetic Connectivity in Protein Families. *Science* **1999**, *286*, 295–299, DOI: 10.1126/science.286.5438.295.

(30) McLaughlin Jr, R. N.; Poelwijk, F. J.; Raman, A.; Gosal, W. S.; Ranganathan, R. The spatial architecture of protein function and adaptation. *Nature* **2012**, *491*, 138–142, DOI: 10.1038/nature11500.

(31) Klyshko, E.; Kim, J. S.-H.; Rauscher, S. LAWS: Local alignment for water sites—Tracking ordered water in simulations. *Biophysical Journal* **2023**, *122*, 2871–2883, DOI: 10.1016/j.bpj.2022.09.012.

(32) Altis, A.; Nguyen, P. H.; Hegger, R.; Stock, G. Dihedral angle principal component analysis of molecular dynamics simulations. *The Journal of Chemical Physics* **2007**, *126*, DOI: 10.1063/1.2746330.

(33) Mueller, R. O.; Cozad, J. B. Standardized Discriminant Coefficients: A Rejoinder. *Journal of Educational Statistics* **1993**, *18*, 108, DOI: 10.2307/1165185.

(34) Pedregosa, F.; Varoquaux, G.; Gramfort, A.; Michel, V.; Thirion, B.; Grisel, O.; Blondel, M.; Prettenhofer, P.; Weiss, R.; Dubourg, V.; Vanderplas, J.; Passos, A.; Cournapeau, D.; Brucher, M.; Perrot, M.; Duchesnay, E. Scikit-learn: Machine Learning in Python. *Journal of Machine Learning Research* **2011**, *12*, 2825–2830.

(35) Chutkow, W. A.; Makielski, J. C.; Nelson, D. J.; Burant, C. F.; Fan, Z. Alternative Splicing of sur2 Exon 17 Regulates Nucleotide Sensitivity of the ATP-sensitive Potassium Channel. *Journal of Biological Chemistry* **1999**, *274*, 13656–13665, DOI: 10.1074/jbc.274.19.13656.

(36) Yang, H.; Yang, M.; Ding, Y.; Liu, Y.; Lou, Z.; Zhou, Z.; Sun, L.; Mo, L.; Ye, S.; Pang, H.; Gao, G. F.; Anand, K.; Bartlam, M.; Hilgenfeld, R.; Rao, Z. The crystal structures of severe acute respiratory syndrome virus main protease and its complex with an inhibitor. *Proceedings of the National Academy of Sciences* **2003**, *100*, 13190–13195, DOI: 10.1073/pnas.1835675100.

(37) Williams, C. J.; Headd, J. J.; Moriarty, N. W.; Prisant, M. G.; Videau, L. L.; Deis, L. N.; Verma, V.; Keedy, D. A.; Hintze, B. J.; Chen, V. B.; Jain, S.; Lewis, S. M.; Arendall, W. B.; Snoeyink, J.; Adams, P. D.; Lovell, S. C.; Richardson, J. S.; Richard-

son, D. C. MolProbity: More and better reference data for improved all-atom structure validation. *Protein Science* **2018**, *27*, 293–315, DOI: 10.1002/pro.3330.

(38) Berendsen, H. J. C.; Postma, J. P. M.; van Gunsteren, W. F.; DiNola, A.; Haak, J. R. Molecular dynamics with coupling to an external bath. *Journal of Chemical Physics* **1984**, *81*, 3684–3690, DOI: 10.1063/1.448118.

(39) Parrinello, M.; Rahman, A. Polymorphic transitions in single crystals: A new molecular dynamics method. *Journal of Applied Physics* **1981**, *52*, 7182–7190, DOI: 10.1063/1.328693.

(40) Huang, Q.; Gruner, S. M.; Kim, C. U.; Mao, Y.; Wu, X.; Szebenyi, D. M. E. Reduction of lattice disorder in protein crystals by high-pressure cryocooling. *Journal of Applied Crystallography* **2016**, *49*, 149–157, DOI: 10.1107/S1600576715023195.

(41) Best, R. B.; Zhu, X.; Shim, J.; Lopes, P. E. M.; Mittal, J.; Feig, M.; MacKerell, A. D. Optimization of the Additive CHARMM All-Atom Protein Force Field Targeting Improved Sampling of the Backbone $\phi$,$\psi$ and Side-Chain $\chi_1$ and $\chi_2$ Dihedral Angles. *Journal of Chemical Theory and Computation* **2012**, *8*, 3257–3273, DOI: 10.1021/ct300400x.

(42) Zhu, X.; Lopes, P. E.; Shim, J.; MacKerell, A. D. Intrinsic Energy Landscapes of Amino Acid Side-Chains. *Journal of Chemical Information and Modeling* **2012**, *52*, 1559–1572, DOI: 10.1021/ci300079j.

(43) Huang, J.; Rauscher, S.; Nawrocki, G.; Ran, T.; Feig, M.; de Groot, B. L.; Grubmüller, H.; MacKerell, A. D. CHARMM36m: an improved force field for folded and intrinsically disordered proteins. *Nature Methods* **2017**, *14*, 71–73, DOI: 10.1038/nmeth.4067.

(44) Mitchell, M. R.; Tlusty, T.; Leibler, S. Strain analysis of protein structures and low dimensionality of mechanical allosteric couplings. *Proceedings of the National*

*Academy of Sciences of the United States of America* **2016**, *113*, E5847–E5855, DOI: 10.1073/pnas.1609462113.

(45) Hekstra, D. R.; White, K. I.; Socolich, M. A.; Henning, R. W.; Šrajer, V.; Ranganathan, R. Electric-field-stimulated protein mechanics. *Nature* **2016**, *540*, 400–405, DOI: 10.1038/nature20571.

(46) da Silva, A. W. S.; Vranken, W. F. ACPYPE - AnteChamber PYthon Parser interfacE. *BMC Research Notes* **2012**, *5*, 367, DOI: 10.1186/1756-0500-5-367.

(47) Kagami, L.; Wilter, A.; Diaz, A.; Vranken, W. The ACPYPE web server for small-molecule MD topology generation. *Bioinformatics* **2023**, *39*, DOI: 10.1093/bioinformatics/btad350.

(48) Wang, J.; Wang, W.; Kollman, P. A.; Case, D. A. Automatic atom type and bond type perception in molecular mechanical calculations. *Journal of Molecular Graphics and Modelling* **2006**, *25*, 247–260, DOI: 10.1016/j.jmgm.2005.12.005.

(49) Nüske, F.; Wu, H.; Prinz, J.-H.; Wehmeyer, C.; Clementi, C.; Noé, F. Markov state models from short non-equilibrium simulations—Analysis and correction of estimation bias. *The Journal of Chemical Physics* **2017**, *146*, DOI: 10.1063/1.4976518.

(50) The PyMOL Molecular Graphics System, Version 2.0 Schrödinger, LLC.
